## Supplementary file 2 for "Hairless as a novel component of the Notch signaling pathway"

**Supplementary file 2.** **Full-length Hairless sequences**

>Anopheles gambiae (mosquito; Diptera)

MYVINPSPASQDLQPTASSYVAARHSESIELGQQQQHKEILSQILTKCNADMSDQVANFSCPDEKTPPRSAAVPVKEEAKSPASSSPAATPAPSSSPCSEVGSGHGAKPTPAVAGSSPKEATSTPAAGEGKTGRRTDGEIRSPLPTLKRNPSLDGSPLSSASSSNAASLSPSSLDGPAKSTPPKSEPMAETVQSLAVASPVSSNFSSVVHPKERALKSIAAAAASNERNNATNGGSSTPAKPASKAVGAVNAGGRAVLNGTGSSSATGTGTTAHGGRLQFFKDGKFILELARAREGEKTSWISVPRKTYWPPTVSSTSANFHKHESSTSLSFSDDNSSIQSSPWQRDHCWKQANPRPNISKELSLYYFRPLKARGHLTHDRRSWCRRKRRRPYDSHLPLEIKSKPQAQKTTTVENGNAKKDDSDHKEAQSATGQEDGETTENLRDKCNGTTPMDTSEETTAAGENNVKKRKGTKQDGPMVEDDDHKQSNGHHPPANGGTHCRPSRCKRPEGKDLASIIKTLVEHQQTKSINAPGASAGAGTAIGPFRLGTIAERGNSTSSTSFRSSVFASSAASQHVSPRKRILRELEKVSLDDAGSSGTSSTKRSRPKAGSAAGTPVIVTATTATPLTTLTTNGTGSPLNGGASSGSNSSSNSSSSNSKTNGHLNGSGSIANGTTGAGGKQPINGAGSGISPVTNPAPPVPAPVSRPFSSYSITSLLGHNTSSSGETSSSANFSPSDAMQRKTDVSVASVTSTASHHLYHHPQHPLSPHHQYQGSLYLGKEPLAVGSKSPIMAESINHAHYQGQLGSTRYGYGAKKRSPSYGGGSGTAASPGATSGGSSVDPCANTIRSPDLSPSPEHHHHQQPQHQHQHQQQQQSHTSAQHSHHGLHHTPGSASSGGFSRYRQHPYGGSPSSYSSAPSSSRFSPSPSTNDSATTPPYSGSGAATGGARSAISYRPGYQLGGSSSQGHSPPTSNGSPQHYSRASPLNFGRGTQLSPPPPLGSQHHNRTAGTSSSPRASPSSGGGGVTTSHASTPTSAMASLTTGTSPGSTIRTVPKKTAALRQPFSGGHSPSSSSPSRERSNSNASSSSHKKDLHPEGSDSVDGASLNHYGTTPSSPAGVIRPNTVIASPTAHHPVPNAFYPLYQAAQLGAPSSSSLMNAAAVAAATSSVPFHPATLTYYQQMYTAATMAAYRTPLWMHYPGLPGVAPHGPPHMIQPHHQAPPPSSVAPNISSADRRLIDRSPPTPPPPPPQPTSGSSITDPAESQRLQHSILATPSSTAALNYSVTALSSSSRDIINGGSAFTSPSGSSWPTDAVGYQHHPDHSARSTAAVSGIASATAKDETSNDVPLNLSKH

>Belgica antarctica (midge; Diptera)

MKAKLKEKFNVHKKAASFTKCVKMSKEVENLKNGILNGESLLSHIQHQQQKQSKLGNSNADRDSKSIVGGRLQFYKDGKFILELARARGEGEKTGWISVPRKTFWPPTISTSSSTTFSKHESSTSLSYSDDNSSIQSSPWQRDHCWKQTQPRHNISKEMILFFHRPERLRLSPQSQKIALAKRRRPLDKLGDIKFIKQEKKEDSDSEDSKSKSDEDKLNGDGEKEVSPDANKAEPAKDIGKLDTILKKLADRITSVLAMNGNSSNGHNGSQTPTTYSMASHIASTINTQPHQHVSPRKRILRELEKVSLEDTKRSRPKATVNGHSIPISSVSPQMTNGNLKSLPHEKAAIAPPVSRPISSYSITSLLAHNNNSTTASSVGSGCSSINDHSNDSSSASHYQQQRLSLSQHSPPKSPSMVQPKRKSPNTTPPGNGNGNSYHSPNHSPSPEHHAFHKYRPTTTTPAALSSSPYGNSYNSPNYMRGSPSPHGDGYNNNRLRTTVSYQHSPSHYDSSPSQSFTAPRDSSLSPNVERSNTSSRSTPTSSGSSVIRTVPKKTAALRQQFSSPTMESANKKPVIKQEKPMEVDTLMRPALVHQQVPHPAQMYPYMYPPMGYMPTAGIPPYYHPAFYNPAMMAAAAAQYRIPGFPPGAMQGYPGAALSPVSAAQMTGASPPITSYEKALNGQRNSLISSPPATHSSVSPYVPTSPWNPIPLTNHSINDGNLIAKAKDEPSSDVPLNLSKH

>Ceratitis capitata (fly; Diptera)

MAAAAALGSSCTTSLSAHLISSAKAPDIDVIDCTMNAMSSVPLNNHSNVYSDKDKVNNKSEKLLLHRNGGGKNNDIEANFSAGSGTFGGRLQFFKNGKVILELARSKDGEKSRWISVPRKIFRAPSATSSTVTPTLATSIAYSKNESLTSLSFSDDNSSIQSSPWQRDHCWKQLSPRSGISKAMSLYYRRPRAASLSKIAVFLARKKRRKPCDKNLCPIPIFEQLSSIVLKTPGNFCSLNTNDIQNTNLNSIVLKTSEKLPLSNKNDIKEINRDNETENDMKNNCSVDIGAALINEVDGCSSSSTATLTGNDFSDNRMEEGPTSLAPSFLLSSCHKENKATVNKESKDVDIDVTVEDIKSNILEKVKTTENENIVSTTTRDEGSEPELNGSSNDVPVLNSKVDKVDPNDQGFCVKLNEDQRLKAPKQRVKLNIIVQKLIDRVPERLAQLSRLSQSVGSSNSIQGHSNLISQKVPQQCTSSSTRLVEYPQQHVSPRKRILREFEKVSLEDNCNVAGGKRSRAKSNASSSSASSYRSMESPNNNRSVVNTTSKISSSGNHTISENNVHRPSPSTYAKSSTTSTLMTTAPTRLYSSYSINSLLGGSGCSSSKAPTLATTNVGTSKRSKDITANPMGYHPKQHQSSSQQHYSDPSYLRAMLATPKSPEQCSGVKSPSIPVPKIRSPPYVSPLRDSQALSEVGTFNNKTRYGSDHPLNLDSSSPMHSQNQSYFLPYTSLSKYVSTISNSSNTTTNALNNSGCTADQHSSRFPSAFNRTSSPASVTNSSNQPASTSIGNSRENSPSNSRNMTDSPRDTVPRTVPKKTASIRRQFASPTTAVVNSSSASSVDRPFEDTYSSQDRITPVSGPNICGGGPHQLHRGSSRGNIVPSPLQHYYMYPSTTPNEASSPVQQSSSLSPGIVLPVVSGGTHSSVSTAAQYIPSIMPSAYMNPYFALAALRQPQLWSHYGSASLPVHMPSITNSPSLRLSPSSFHSFAYNGVNAAMAAVMQHQQQQILASSALMHGHSLSRTPVMVAPHNTINGSSGSQDSEAILTEQTTDLAADEIPNNATGLRLETGRCSSIKEEHNSDVPLNLSKH

>Lucilia cuprina (fly; Diptera)

MKILQNETTNSSNSTTTSPNSKEKQQLPTQSSSLPQQPQQNFVEKRKAKTLKNKILQNVAQQQQQQQEQEQQQQNANNNVNNNKHHTPSAMTDERTTTTNNSNNNTKTNKSNNTNDSKTTRATTTNLLEDSLKDSSLGTSNNNNDNNNKNSDLIIAEAAKILLKNGLNGTNAAAAAAVATATATLLTAAANFHQLPTVTNSLNSSSTSSSNSNLSTSVSSTSSSCPSSTVSDTSVLAANVTTAAAAAHAQLVAIAAAAAAVNATNHKRLAAAPTSSASSTSTTSSSSSALSSCKSPKQTINNGLLANATGIQTATATPATAAHFQKPYKTSSCRSTMSSAMDTGIIDYTINSNSNTSSSSSSTLSPSAALSSVINLSASANSSSNNSNSAHHKLHNNSNHSHSNRSDIGRPPISHNHQTSNANNHKNTVPGGGVGGLGGRLQFFKDGKFILELARSKDGEKGGWVSVPRKFFRIPATSSTVTPSATASTSAAAIPSALASSPAAIVGGASYPKNECSNSLSFSDDNSSLQSSPWQRDHCWKQAAPRKNVSKELTLFYRRPAYHKLTRQALLLSHRKRRKPYDTTKTLKNDLIFVLNSKKQLQQQQQVDKSTISNKKEVELKENIDKTSEKGADENKNEKVDKGGDAKCIKSCTEDVKNEIDDGENKNNGAKEDFDKSSDIKAIIKTENEESAVETEEDVDVEMTKLSSPCKSTENEANIKNVEVIKENDQVEKMNTSDYNDDAEEKPKPLNQIENSTALNGNVLRNDDLLEKQQQQVTDTPGEGAKSKTGDEEKRLKKHEHYKTRAKLTTIVQKLMDASSSRLAASTHLFKTSNQQTTTTAATPTSSSSSSSNSIPSTASASAGLKNQNSKVSPPPNTAASRLVEYHQQHVSPRKRILREFEKVSLEDNATGAAGGKRSRAKGNSNTINTTMAGNNSNIASSSSTSTTSNKSMSFNAHNAKSASSTASSTAPTKLYSSYSIHSLLGGGNATSTSSSTPAVKKSTESPATITSSAMAYSPYHSASFPSNLHSNNSPKSPDGHNICAGGGKSPSNSATKSKRSPPYSSPMRETQSRSPVNSGGYADYSGRRSPRSTPDTAHNTFNKMRYAGGSGGVTDTAASSSNHHVQHHQQQAHHLQQHQQHYHHQPQTYYSPYMMSPHYVPPLSGPSALSPNASSANASTSSTTSSRSPRHHPSAFRATTPTSASALSTTHNIPNTQKGDLSPQRSTTASPRETTPRTVPKKTASIRRQFASPTSATQNHNNNSSNCSSPTTIIDNRSEDQMTNDRRTPVNTTTMHQQHQQPQHLMQRGSPGNPSLAGSPVHPYSYLYAQPPNGGNSPHSSNAAPSNHPSTAMSPSAVNASAAAAAASYIPSVVGLPYYHPYISTLAAMRHPQMWIQHYQSAAAAAANPAALMQSRHPAMLSAAAAAGAAARLSPPYHGFQYNGVGNAAALAAAAAAAAGFGSTATPSHHPSSALSHSNSMFNPMANSSLHGIQHPVTAMQTAAAGLVPVSAAAAALSAAASSSSPITDLSSKSSSKNDYSSSANTTTTLSSLSTGTEGGGGGRSGLGGGASVTESSGNSMYTISNKDEQSSDVPLNLSKH

>Mayetiola destructor (fly; Diptera)

MTDEHGMLKNGSVDSVGIVRRETTTIATIGNNSTQTNNINNDTNNNTNDNNHDNSTRNNGNERESNPQSPIDATNGNSIDAIVSGNCNKQPNNNTNNYSNNGSPASNPVGGRLQFFKDGKFILELARSKDSDRGWISVPRKTYWPPAALSQNNVNNHKNESSTSLSFSDDNSSIQSSPWQPERSLKQPSPRKNISKDLNLCYCRPVSIVLNGKSKNISKRIRRQPHSKKYENICVEYAKRAGENVVESLADDKKPDTKMECKNELDEDKENEANGQMETEDTVDGSVPVKDERMPSGEAEQMEIKASESNSNDAEPNNDIKIDSRHNDDDDDDSTKNETKHKCFKKRLIRSAKIGTIIEKLITKVSEQPVNVSTTQNGPGFHLTSSSRSNDLLYPHQHVSPRKRILREFEKVSLEDRNSTNLKRSRSKSNASSDSPNRTVVTQHTVSIANSTKDDHSRFTNGNQIPLKRSSPTDVSRSESAFKHSKNPSTISTTINSQTIITTNENPPQTISKPLSNYSIISLLGHNNTTSDKYENNDDQKPSTNDAGSPRSPHSYNHQTLASNRTSFLAKKRSPTNNSANIHSPVNYRNSPRSPNNNSPSPGLSQNSHSHRFQHNSLPASISSPTSGFHPYLASARVSPLSSGTLSPPDMYRHRSYRPACSPSTISNSSGGLSHSYASLNSSPTAFANRYSPSTYSNSSTSLKTSPTQSSHSQNLSSAFNVSNLLPQPTNDGSSNNDRLSHNYSPAKNSIGTTNSAQNSPSNTTISATTIPKKTASIRQKYGSVSPNGIMNANHTEQTKNDGPAARKLHNDNNAYEAKASTKRSRSPTDFIKKHQQQEYEKDLQARYNAELARQQSFMSAMQQPPVGNPFYNIYHAAAYAQCTGINPYYHPNDLLRKTHPSELIHKNLPPHHSWIDPYTASLMHYSGVGPGLVSDKNSIPKPAGGELHSMLMSPYGLAGSSAAHGLTSAADVEMWKTNEKKAPISPTTSAFRRQSQPSYNDCEPISLIKDEQSSDVPLNLSKH

>Ctenocephalides felis (flea; Siphonaptera)

MRIQDCEIIPTKCANMTEESDFKYGMNGGSPLKQDNTDHKTIASPDRAPTDKLSQVCTGGGRLKFFKDGKFILELERARAGWVSVPRKTFWPPPAPANIPGSRLEGSTSLSVSDDNSSIQSSPWQRDHCWKQTSPQRSSSQETSFLYKRLKGIRFAHHLSFRKRRRPFDTSMNDAKCYTSNGISSNPSNRIFRKPGILANIIQRLWDRNGASIKQIKQTVSIDQHAAMVSPRKRILRELERVTLDDLYNKRRAKVAQATSTAQVAATSASTGVVHQNGSVAATTAVTTMAPTQTSSKTCSYSIDSLLGNNSTDTTDSNKNKSSVTHQRYQAAPLPVHFSTYYNPLGHMYPTYPPRYAPLWSQYHPSLMNSYAGTIAPPIWAHSSTQHRNAPAAGVPPPAHAETSNVEEDVPLNLSKNSR

>Calephelis virginiensis (butterfly; Lepidoptera)

MHIQEGTNTTLTTCTKMTEEVDRRHGVNGTSSSSKGGNEDPPVHSSAPSATGGRLQFFKDGKFILELVRGCAREGERAGWVSVPRKTFWPPAAAPLASPLAPPPSATSLSDDNSSLHSSPCTSHRDHCWKQPTPRRNLSKELAMYYCRPTTLRNDHIHAASRHKRRRPYDTSYNDTIPNGTVRKTPTNGLCEKKRNGDIGLTGNKHCKDEDTVDGPSEIKQKTDDERWQDFDREKYYHMKLKKPYQYHKLRVYKKTKSPMRRKELTRVIERLRDKVATFPVQLVNAKIANCKQEHTIVSPRKRILREMERVSLEEGTKRRAKTVPALSTASYPPSPGPSSARAPAAAANGAAGRREAARGVSSYSIHSLLSMDERAPPTPDPSPSPPYRLPEPFRAAYAPPPRWGPPPAPAHYALYPYPPVFRPAPLWMHYAPWPPLAHPVLTEHIHKDEPTTDLPLNLSKH

>Pieris rapae (butterfly; Lepidoptera)

MHIQEGTNTTLTTCTKMTEELDRRHGVNGSKGLNDDPPAQNATPAATGGRLKFFKDGKFILELVRGGERSGWVSVPRKTFWPPAGAPGTPPPPHAAAPCATLSLSDDNSSLHSSPCASHRDHAFKQPAPRRNLSKELAMYYCRPSTLSVALVRAASRLKRRRPHDPVHHDASFEEAKRNGDVADDKTCRGDDVVDAPIETKTKHDNDRWTDLDREKYYHMKLKKPYQYHKLRVYKKRKPIAKRKELSQIVDKLREKVAALPVLVNAKLANCRQEHVIVSPRKRILREMERVSLEDQATKRRAKTVPTLSTASYPPSPGPSHTPHRAHEPTRAQNGTLPKKETVSKNVSSYSIHSLLSMPEETGTRRSPEAKRSPHSYPPSLKTESPSSVTSPDLSPSPDGYRYRYGLSLDSPGRGAARDSPTPPAGAGVAYRGYAAPGYARAPPTYRGSPGREDWPPYVYGYAYPPLYRAPAAPAPLWMHYAMSPGVPPGAWPPLAHPLLTDHIPKDEPTSDLPLNLSKH

>Spodoptera litura (moth; Lepidoptera)

MHIQEGTNTTLTTCTKMTEEVDRRHGVNGNSSSKGASEDAPVNSSALTATGGRLKFFKDGKFILELVRGCAREGERAGWVSVPRKTFWPPAAAPPTPPAAAHPPSASLSVSDDNSSVQSSPWQRDHCWKQTAPRRNISKELAMYYCRPSTLHTAAIEVASRLKRRRPYDTTHLDGIVNGTNSRKTLTNGVNDKKNGDVGSPDNRACRSDDVVDGTGPKVKTNSDDDRWTDYDREKYYHMKLKKPYQYRKLRVYKKTKPALRRKELTKTINKLRDRISSLPVPVNAKLANCRQEHMMVSPRKRILRELERVSLEDQSMKRRAKTVPALSTASYPPSPGPSHTTHKPVVETPIRSHNGTAAPRKETPSHVSKNVSSYSIHSLLSMPDDSPTRRSPEAKRSPHSHPPSLKTESPSSVNSPDLSPSPDNYRYRYSALMGSPGQGGLARDSPTPPLELARYRAPYPPPTSPYGVRQWGGVSVGAQYRGSPARDEWGRDMAYMYPYSYMGPHLYRTPPPPLWMQYPMSPGVPPGPWAPLAMPLTNDHIPKDEPTSDLPLNLSKH

>Trichoplusia ni (moth; Lepidoptera)

MHIQEGTNTTLTTCTKMTEEVDRRHGVNGNSSSKGTSEDVPVNSSALTATGGRLKFFKDGKFILELVRGCAREGERAGWVSVPRKTFWPPAAAPPTPPAAAHPPSASLSVSDDNSSVQSSPWQRDHCWKQTAPRRNISKELAMYYCRPSTLHTAAIEVASRLKRRRPYDTTHLDGVVNGTSSKRTLTNGVCDKKNGDVGSPENRSSRSEDVVDGAPKAKTNSDDDRWTEYDREKYYHMKLKKPYQYRKLRVYKKTRPPIRRKELTKTINKLRERISSLPAPVNAKLANCRQEHMMVSPRKRILRELERVSLEDQGMKRRAKTVPALSTASYPPSPGPSHTAHKPAVETPARSHNGNVPRKEQPVSKNVSSYSIHSLLSMPDDSPTRRSPEPKRSPHSHPPSLKTESPSSVNSPDLSPSPDNYRYRYSALMGSPGPGGLARDSPTPPPELARYRGYGAPASPYGVRAWGGAAPAAQYRGSPARDDWARDVPYMYPYSYVGPHLYRSAPPLWLQYPMSPGVPPGPWAPLAMPLANDHIPKDEPTSDLPLNLSKH

>Limnephilus lunatus (caddisfly; Trichoptera)

MHIQERNNTTLTTCTKMTEEVDRRHGMNGNSSSKGNSDDSSVNSSAPPNSATSAGGRLKFFKDGKFILELARAREGERVGWVSVPRKTFWPPTPSGPPPAPSTPSAHSTLSALSTLSARQERATSLSVSDDNSSVQSSPWQRDHCWKQTTPRRGLSREMCLLYLRPSCLSSRLLRCPAARLKRRRPFDPTPVPPVPPQPTNGSPSLPASASPNPADSPHPFRGTDKPTAQGKFKFFRNKSDNAIEPVALRVYKKTAKKAKRREMIAIIDSLKERLPQFTPVNAKISALNCKQEHSMVSPRKRILRELERVSLEDQANKRSRARHTPPATAPQPGPSCAPDPPKLYNGSSHSHSRPHPHPPSKPVSSYSIHSILSMSEDSAPGGSPDPHSKKPHSYSSVKTESPVSAHSPDSLSVKKPHAYACAKLDSPVSVHSPDPSPSPEHYRRAAGVWGRSPPPVRPYMPPRAPPELTDYYYGEGYSLYPGAYLAPSPYLGSYPGYVPARAPPVWLQYPPQLPPGPWAPLLSDHTPREDTSDLPLNLSKH

>Harmonia axyridis (beetle; Coleoptera)

MHISESRECPNTILTKCAKMNENSHKKLSINGNSDGTVVKVKEENPKNYYVPPTSGGRLKFFKDGKFILELERAKEGERVSWVSVPRKTYWPPQGAATTVTCKQEGSASLSISDDNSSIQSSPWQRDHFWKQTNPRRNMSKGMSFYYWHSKNTPRLPRSKKTRRPYSLEPERETSNSACGDQKTELKKCNKFENYKKRRSLLSVVQDLIDKNVSKSPPRIETVVSPRKRFLREMEKDKNSLEDSNLKRSKNKMTSNQASLSVTMSSPLRVNGHSEDKPVASVAPSTVGKNCSYSITSLLSDDRVPVKRSPSNSPSHFVPVTTNQMNYSMASVKAEDLWYSESVERFRSIELSHADKNSFHSFPHPSYIPPFVYPYSFPPCYGYNRGFPPIPQGMYHHPTAHHLPVPMMRQEAPSCSWKTERMEEGEVKKEDNNLSDMPLNLSKHAG

>Pogonus chalceus (beetle; Coleoptera)

MRITNSHRDHTTNLAKCTKMTEGEHRIHGMNGVTDSGTAAVQEETTQSTPPVSDTYNNLGGRLKFFKNGRFILELARAREGEKVAWVPVPKKTFWPPQGSATSLSTYRQESSTSLSVSDDNSSIQSSPWQRDHTWKQTSPQRNNSTNLDFYFYNNKRKNKNRALLQKSCWQYKLQRAPYLIITEQQRTKCITLFNNSGNKCNNSKCINNSNNKSSAISSNIKSRQTLANVVQLLWNKSLQALNNLTSPRKRILRELERVTIDDLSIYNKRCRGKVNQQSKLLTTTPHQPQQLQQQQQQQHQLQCTTNNNNNNNDNISTNVCRSDQTDINGFDYHRVVSSSSSPSPSSSSVINPCVSVTSTKNSSYSITSLLGGIIETNRDGNNQIEKLPSSQYTTKITTTPADNHSHINTTYSSNNTYISTDEVDGPSSKNTLDNILLKNKRSPIIRGGTSPSNINNSTIPSPSSHYNYSAVNAAHHQHQIPPSHYMSSPYSIYSSVLHQHHHQHLAAANYISPYYHHPQSSVGAAAAYVAARGAPPPGYYHHPLVSQTLNPTPWNTYEQHQGNAVHRDDSDVPLNLSKHAD

>Pyrocoelia pectoralis (firefly; Coleoptera)

MRPRGRSPLTKCYEMRIVDSREASTISSWTKCSKMMEEGHRIHGINGNTDGGAVVPKEETSKNAYNQGGRLKFFKDGKFILELERAQDGERVSWVSVPRKTYWPPQGNAPSIPNYRQESSASLSVSDDNSSIQSSPWQRDHSWKQATPRRNQSKEMTFFYYHLKKFPIWRTCKNRRRPYAISAVDKTYLTIKSERTEQNSVCNKVSNKLMERSSLMTVIQMLIDKSASSTPPRSEIVVSPRKRFLREMEKDKLQLDDACQKRSRNKSAVSGSAPPVTIKVESPIRLNGITSSITEDMRAPRNCSYSITSLLAEDRSRCSPTDSPSHFSPVPQYCSPHEDPWYSESVDRLRSIELSQVEKRTLAAYSSYPYVSPYMYPYSLPPYYSSGIYGRNYMVPPIYPTPPMPVSMRHDTPSCSWSADTSKDLDHRDDTIADMPLNLSKHAG

>Sitophilus oryzae (weevil; Coleoptera)

MHISEPHTAPNSAHILTKCSKMTEESHKRHGMNGNVDRISKEQCHKTNYGQGGRLKFFKDGKFILELERAREGERVSWVSVPRKTFWPPQGTASSTPAYKQESSTSLSVSDDNSSIQSSPWQRDHSWKQTTPRRNVSQEMALYFWRPKRRRRQCRGRGVKRRRPLSVVLDVKEEDAAVAAGRVREKRRGKSLSLIVQSLLERTARTSTPPRPETVVSPRKRFLREMEKERTPSASSSPSSDRPGSSDDGSTNSAQKRSRGRPLGSGGPPVAKTPPASAIRGEKLGNSTPPPLQTSVRCNGVEDVSMNSSKPAKNCSYSITSLLAEDHHKSSAKQSPNRSPNRFSPVVTQMGVPQATRYCSPVTSEDRLYSESVDRLRSIELSQVEKCGYPASYPPPPQPYLTTPYIYPFPPIPPYYGPGVYGRAPAGYVVPQPLYHSPHHAHSPTPHPLAAPHHVAAAHHAELRRERGGSTAPAWTNSGGGEEQRGRREVKDAREENVDMPLNLSKHSG

>Mengenilla moldrzyki (parasite; Strepsiptera)

MHFAIKGDVMGQETESDSKSNPDSVKSLIATDDNNEQSDNKRGRLKFFKDGKMILELTQARLGEKLAWIKVSNGTFWPPQGVVAPVLKADIFTSDDNSSLQSSPWQRDHTWKQIRPKKWSNDILVSFCYMKSRHMKLHQWIGEKSLSRNPFRVQNNWWPVKCRKCSSKVENEIDFRTNVCNNNTFNSGNSNSAAKTKRSLHDIVRDLSLRNQAQVVINRNMCMCSGGELLMPSTTASPSPRKRILKDLERVSLLEDVTHTINGFGNGNNGHVCDTSELGNINGNSSGSGNGNGSGIGVCEDRNVIVDKNEKCKKRARCIGTKQVPSFKTDKKTVPKNNSYSIMSLLSRNDSNELSVSTNTSPPSLNSSPATMLNPSVVSYYPEQSLLQSYLTTPPVAHQRNSSFYSTYSRTGLFYSYHNRSLPLSANNLSTVQNSNSPSCSWFRDEYVSDVPLNLSKHAG

>Stylops melittae (parasite; Strepsiptera)

MHIALKNHTANHVDHMIANCKQIGDETGPNDQSNGDNNDDSVDNKGGRLKFFKDGKMILELTQAKFGEKLSWIKVTNGTFWPPHGLITAIMRHENCTSDDNSSLQSSPWQRDHSWKQIRPKKCPNSMKNMEFFYKKPKNKKSHRWLLNCSKKKMSRNACQEIFVLKSNMCKKCKVHVNDCGTNKCNTHLSSLSSSFGATNGASKSLDLTIRKLMLRFHSQSVIHKNICSGGEVLSPTTNTTSRKRNLKDYDKVLFLDESNGNGVPITTANGSSVNGNGGNSKKRARCVGTKLVPTSNKTEEKTLPKSTNYSILSLLSKKDSNESIQGHDKHHRNVSTNTLSPSSLSSSPTSSSSSTSVPVPVPASTSVPTLTSVSATAAVTTTAPLPTVKTEPASYYREYSLLQSYLLTNPTPPIAHQYNGQIYLNARNRPDLFYAHHNNSFTFKHGFNFNHIHRNLPLSSGSQMHINDINPLSSDVPLNLSKHAG

>Chrysopa pallens (lacewing; Neuroptera)

MHIQESHQTQENKQAKCSKMTEDGHKKSGMNGIIDGGSGNVGAAIIKNNSRNDTEHRATGTASSTGGRLKFFKDGKFILELARAREGERVSWVSVPRKTFWPPQASTGSTPPTHRQESSTSLSVSDDNSSVQSSPWQRDHSWKQNSPRCNVSSEMTFFYHRPSKLHVPVVRHKRRRPFDSSSNINNIRTSSEHGERDECDSSVQSNKIVKVNNTEDIKKDEGVLSGVGIGGAEGTNVNDNNSNNNSDSNKIKGVNKKRQMRKLVDIVKLLWDKKPQVNNYSTFKSVLMSNIRHEQMVSPRKRILRELERVTIDDLSNKRSRAKPAQSITFSTATTGSSPSRAHNGTAETSSSKNTSSYSITSLLGHKKEEEVSNPTPSISHTEFHNHSTQLSPNVSQRVLLNNVSVSPKNINYSHSPKIIAHHDSPDLSPSPEHYRYVSHQNIAHSSPTHALSPPPDVHKLYRVHGSPNQYNSLHGLVLHQKSPQYYTPSHRSAAASPCSPRSPLDDSRSREHRFIEPSSEKIDNVGHRVLEMEHSLRPTFISPYLYSPGPSPYISHPAYYNAGPYRSPYQAPYPGESLTHSPSSQPRNYATHQNWSQVNQNHALDALRDDIASDVPLNLSKHAG

>Conwentzia psociformis (lacewing; Neuroptera)

MRVENVSDTVLQTRAKCTSYPTMTDDSSATGMNGIIDATNVVAPNKQQPLIKNETPLTTGGRLKFYKDGRFILELARAREGDRMAWFPVPRKTMWPSPATISTGTTPSTPRQESSTSLSVSDDNSSIQSSPWQRDHSWKQTTPRCNVSSEMNFFYSKPLRLKGHTIKLFNKLLKRRPFYNFHDRGNCNATNTCRTSGIVNLSKPKPLSSQSPAPNPPSNEKTVELPDMPDINVETTTNSSNSDSLTNKTVQSQFDSVNNNGKAPVNSRATKMKLSDIVRMLIDKRTSCLGAFNNFTASSSAPVRETSMSTTNSTSKVINSSHSMSAWSETSPSGVPLMATGTNPTSSSASNCKPFKTHLLRYEHQMVSPRKRILRELERGSAATNDSDNFKPPKFKSTNIGPMDKMHNGATEGQSVTASAKGTTPKSSSNYSINSLLGNVKTEKPDDTPDYRKSKSIHLAYDAQNLSYARNEQDPSISPPSHIDRYTTPSTSKSGGSPILERSDRDRYVAANSERNTTTPSPPDNYKYMKIKKMAANKVSSVSPSALSTVSRRSPQSPYHSSSRTHSPRHVSSSGSSSKHSSPTDPGSSSHSTLPSPTSGYLPIFGLQQRLTLQSPSSSPDRVLAQHSTLISHQYQREPETEHSPRSVSSLTKTGNATSSSNFGFVPAPFAYPPSPTPPSHFVNHPGYLVYRNPYFPPYSLPGEASLLSHLSEQQLRSQINPLPHPGAMWNHSTRVAAAHNASHTLEAVQEDSSDVPLNLSKNAS

>Corydalus cornutus (dobsonfly; Megaloptera)

MRIQELCQLQTPKNREQTKCSTMTEEGHTRIGMNGSIDGGGSSTVGATGAQSDSEQHGPTASGGRLKFFKDGKFILELARTREGERNMWVSVPRKTFWHLPTNASTIPSSHRQESSTSLSVSDDNSSVQSSPWQRDHSWKQSTPRCNVSSEMTFFYRRPPTHLRHKSRSILRKRRRPFDNSNSMLRNISEGDVTSRLDGSVQSENAATAHPVAVEAEEESEKKFENGQTQERKVVPSAKERRKLSDIIRILSEKKKHISNNTVPGLKAVISSSRHEQMVSPRKRILRELERVTIDDLSNKRSRAKPVSTQQSANLASGSSPRRAHNGSTESTPAQTKNSSSYSITSLLGHRVRDDEPPSPGPPSEYNRSSTQMSPGIQHATVNNLPSPKNVTNYSSKISSHYNDQQDLRSSPEYCRFISQHQSTMVHGSPTHNLSTSPDVCKTGAPPTYHRSHESPNHIRHHHHRQQQNLFSSPLQGLAGHQKSPQYYNPSNSHGCAPASPHSADERFLSPSDKTERSALRRSSPKGFLENKSPPKSFTETIPSGIGPPSAGYACVPPYLYHQTGAAYVPLPGAYYGTTYRPYVMPYTLDPATRPSSPHYRPPIVAAHAWTSLSQVHAQDTLRDDISSDMPLNLSKHA

>Inocellia crassicornis (snakefly; Raphidioptera)

MKIVLRPFGCSVEIIDYEMHIQESQPPPNNSQQSKCSKMTEEGHHNRTGMNGIIDEGNAGAVSSEKNHQTSTATSVCPTSSGGGGRLKFYKDGKFILELARAREGERVAWVSVPRKTFWPPAATTNSVPGLRQESSTSLSVSDDNSSVQSSPWQRDHSWKQNTPRCNVSVEMTFFYHRPSSLQIRVAWRKRRRPFDPTSDSVRRERAAAENLKVKSEAADNEDNISDKIKSEGWDECDNVIVQSSIIKKEINEVSEHCSIKVNTNCVLKLDSKDGIINGSVESQCSNNKLSVVKKTRKKLEDVISALKNKKPSHVPNANLISSVFSATTTIKSFPTTRHEQMVSPRKRILRELERVTIDDLSNKRSRAKPAVTVNTTQTVSFSSTSHCSSPSRAHNGSADDGTAIKTSQNLSTKNTSSYSITSLLGHTKKSDDNSPAHANIYEYSPHSAQISPETRKPMPNNLPSSSSPSPKQHYTSKMIHANTPSHNYDSPDMSPSPEHYKYMASQPVVRSSPTNHPLSPPPELYKSYRLHASPNHFSPPVQHVAGMSLHHQKTSPQYYTSPPGHKGASQPSPVSSSPHSPGPEYVNAPTSYEERYRKFVVDRRQRGSPSLDSDKRMSEPNIEMMQRHLIEVDHASLRTAYGPYIAPYVYPPIHVHPYINRAGYYDPVYRNPYELLPTYPFEALPPRPPSPSHFRVPHSTWGSPIAAHSSALPSIHNAHRKDDTSDVPLNLSKNAS

>Cerapachys biroi (ant; Hymenoptera)

MREKSAECHAHAGHVRGTTEMCSRDPPTLHPTLAKANGKNSSPQSAHHTEGRIDNNLLPTPGGRLKFFKDGKFILELSHRRDGEKTTWFPVPKKTFWPPTSTIPNRQESSTSLSVSDDNSSVQSSPWQRDHCWKQTHPRRRITTEFNFYYRRNPKIPLCVHPRLIARKRRRPLDSTTAVLVSESTVIYPTRASRKLNGASSSGKVLSLIIDKLARLLDPNVVSPRKRILRELERVSLEDQASKRRATPQPVCTTTSAPSTPSPKQLSSYSITSILGEDKPSAEPGFLRTLLKPDDRQPVKYSSSNTYPRVRIDPYIGSSNTTVHHPLYVPTLPPGPYRTPLWMVPHSYPSPVHYPPPMQLYAPPHSHPSSPHYKDYREQTLTPPSDMPLNLSKHAG

>Ceratina calcarata (bee; Hymenoptera)

MREKSAECHAHAAHVRGTTEMCSRDPPTLHSTLTKANGKNSSPQSTHHSETRIDNNLLPTPGGRLKFFKDGKFILELSHRRDGERTTWFPVPKKTFWPPASTTPNRQESSTSLSVSDDNSSVQSSPWQRDHCWKQANPRRRISTEFNFYYYRNPKNRLCAHPRLIARKRRRPLDPTSQLFVSETVTNYSKPARKANGAPTGKVLNMIIDKLARLLDPNVVSPRKRILRELERVSLEDQASKRRATPQPVCTTTAPTSSPKQLSSYSITSILGEDKPSHEQGFLRNLLKPDDRQTVKYSSSNSYPRVRMDPYIGTVNAPLQHPLYVPMLPPGPYRAPLWMHYPSPVHYPPPMPLYAPPPPHPPSPPVHHYKDYREQTLTPPSDMPLNLSKHAG

>Nasonia vitripennis (wasp; Hymenoptera)

MREKSSECHTQASHVRAMTEMCSREPPPAVLQHHHPALGSSKANGKNPTSPPVQQQQHHNHAAHHSQSSPRPSSLVASPASTPSGHEDDASNGASNAAPGNSNNHLPTPGGRLKFFKDGKFILELSHRRDGERTTWFPVPKKTFWPPANVTPSRQESSASLSVSDDNSSIQSSPWQRDHCWKQTSPRRNISTEFNFYYRRNPKTRLSAHPRLIARKRRQPLDTCSSLTQLQESLANHQQQKQPQPQSSTATLANARKLNGAAAAAVPPNKSLSIIIDKLARLLDPNVVSPRKRILRELERVSLEDQASKRRATPPQPANTPQQSNETPAPVSKQLSSYSITSILGEDKPSAENEPGFLRNLLKPQERRHPPQPSQHSPQHSRSPQHHPHSPQQHPHSPQHHPHSPQHQPVYPRSTSRLDPAVAAAYLSPNAAASSIHQPLYGLPMMPPTGYRAPPFCWMHYSQPVHYAPPMPLYAAPPQPPHHTPSPSPPAHRYKDYREQTLTPPSDMPLNLSKHAG

>Synergus japonicus (wasp; Hymenoptera)

MREKSECQTHAGHVRGTTEMCSRDPPVLHPALAKANGKNSSPQPTHHNDSRTENNVLPTPGGRLKFFKDGKFILELSHRRDGERTTWFPVPKKTFWPPASTTPNRQESSTSLSVSDDNSSVQSSPWQRDHCWKQTHPRRNVSSEFNFYYRRHPRNRLCAHPRLVARKRRQPFDPGTLQLGPETASGFSKSLFRKANGAASGKGLSIIIEKLSRLLDPNVVSPRKRILRELERVTLEDQASKRRATPQPINTTTSAPPTPSPKQLSSYSITSILGEDKPSSEPGFLRTLLKPEDRQTMKYSSSQPYPRARIDPYVASANTSIHHPLYGVPMLPPGPYRAPLWVHYSPPVHYPPPMPLYAPPPPHPPSPPVHHYKDYREQTLTPPSDMPLNLSKHAG

>Ectopsocus briggsi (louse; Psocodea)

MLLTWRKTSNYRMRVVEPAVPASCAIMSEEKSLNGSEVGSSIRKPADLTHSPNSEAGETSGTTGGRLKFFKDGKFIFELTHRKEGEKDSWIPVTKKSCWHPVVAHSTASLSASDDTSSVQLSPWQRDHCWKQSAPRRLRGAQLAFTMTPLAPALHARDVHRSLLYRLRRRPWDSFALPSTSVNGWTKPPSRQKRSKGSPGRLSGVVQRLWDIVKSSPSLFSPRKRILRELERVSLVAEEPVKRQRPRAPPAGHSKGVSSYSITSLLGPREEEPSFLRNLLRSPSPKSRQHSPLAEAVVAPAVPGPPSPYVYYPFQSGLPPPPPSYFVAGSTSPPLWTPYPLSSLPRGAGGPYAGLVPSYSSPHFQYPWPGSPDCELKREDTSSDVPLNLSKHGGS

>Pediculus humanus corporis (louse; Psocodea)

MRVVEESASSTSCAIMNEEKSINGTETDDCKSNKSQSTEQNHSPNSDYSEQVYASSGGRLKFFKDGKFILELSHRRDGEKMSWIPVPKKTYWPPPVTTSMVTNSIRQESTTSLSVSDDNSSVQLSPWQRDHCWKQMSPRKAASHDMSFFMIPLPRAFHVRRTNSLTIKRKRRRPFDSIEKVINWKSLSNLNGYQQAVKLKNCAIKNGRLNLLIQRLWDYCFKSSESAQSLQKLSPSMFSPRKRILREFERVSLVGGNLDDQSQLNLKRQRCKTSFENGQSPPTQGNSKGVSSYSITSLLAVKEESQDQESSSFLRTLLKSPKEEPSPEPSPKSRRNSPSQSMQSSPPLPVPSSGDASLRNVQFPSVQFPYLHSPLLYPHFLPQSPAAHHSYFGMPSSFRGASSALWGVPYHHHPHVSSSLPGGSYHGLVSPYPTSQYSYWPGQPPINEIKREDSTSDVPLNLSKNAG

>Bemisia tabaci (whitefly; Hemiptera)

MKAMLIEDCVMSTCATMTEDRSINGDININSKKSTILNSNGPFSPVSNSNIDPDNSASFGGGRLKFFKDGKFILELSHQKDGERTCWVPVAKKTFWPTVGTPRLENCTSFSVSDDNSSIQSSPWQRDHSWKQSNPRRNVSKELQFMVRIRPLPRKYFSSYSVRRKRRRPYDPTKIEVKDDSSHHATSKSVKKEKKEKNGIGAKLSSIVYRLKERIHVSVVKTENTINPARIDPGIVSPRKRILREFEKVSLDEIAVSTKRHRARTAPAPAKPVSSHSITSILAREDEPSFLRSLLGCSGSEASSASFDSSQSRNPYVVNPSPSPESVRPSACSSTTSVTPSPPPPKAGTLVQPQPTHPQFPYAMSMHPPFLPHPQASFYSGLSHYRSSPSYWPLYAMSSLPRNTLYPPMIPAFSPLSPSSWTPLTQPAIGEFKRDDGSSDVPLNLSKNAS

>Cimex lectularius (bed bug; Hemiptera)

MRIKEKSSQETSTCAKMTDEIMNGESKTTNRQQPQQQQQQQQPQPQQQQQNQLSNFQQNNSSSPNRLDFDQTPLCGTKLKANKGTPVGTNHSLKETEKEDITNPDSPNVCNEEYSSGGGGRLKFFKDGKFILELSHRKDGEKTCWIPVPKKTYWPVIGTPRQESSTSLSVSDDNSSVQSSPWQRDHCWKQTTPRPNLSRELEFQLIRPKRFMKFLYTSQALRSKRRRPFDMSKVDETLLQERVSYKPRKRSPLRNIVQSLWERVVTTCKPEPGIVSPRKRILRELERVTLEEATKRQRARAPTRNISSHSISSILAKEDESVLRTLLRSPSPDAAIRSRVPTSSYIPPASHIVHPPLYPASSTTYYPTVTNQYTATHVWPVHYPVSSPPLHLPLYPVPPYSSTWPHLHRTMPQETSPDVPLNLSKNAG

>Ferrisia virgata (scale insect; Hemiptera)

MRIQDREAVNAPCVGTMEAKDVNGASIPATEIKTDLKTEINDASHSNGILSPPTTPSTTTPSESDYAPNGPGSGRFKFFKDGKVILELWHHKNGERMFWIPVSKNSTFWPPTGHCTPRQESSTSLSVSDDNSSIQSSPWQRDHCWKQTVPRKNVNRGLEFSWVARKRKMRFSSLAIRRKRRRPYDPVKFEVDEIFKEDNSSRKFNSNIKRKLIDIIQKLSERRSYENATNSCRTDPGIVSPRKRILKEFEKVSLEEMGNTSKKHRARTIASSASANTVSVFTGSVSSPIANASVTAITSKPANNHSITSILSRDEEPSFLRNLLKSPNESSSESKSSFTAESPSNADIRIPPSRPSHCSTPISASLSPPYPKNLHCPQPIPSYIPPAYLYHSTQPFLSSPPISNHPPYYSAFTSTYRDPSVWSMPSVSSLPRQSVYPAANVPSYPPISLSPYIPVAHSSFQSYSVTENGNDAPLNLSKH

>Homalodisca vitripennis (leafhopper; Hemiptera)

MRIQELGPTPCSKMTEERSRPMNGDLSSGIKKELTCKDKNGPTSPSVTTNCESDSVNSISGGRLKFFKDGKFILELSHRKEGERTCWIPVPKKTFWPAVGTPKQECSTSLSVSDDNSSVQSSPWQRDHCWKQNAPVHNAGRGMEFMMTKRRSHHRLKYTPKSVRRKRRRPYDTSVVVMFADWNEKPPIKSTPSRPVLGINSVIQNLWERVVRADPGIVSPRKRILRELERVTLEDQNNSKRQRARPSPEQPKPVSSHSINSILAREDEPSFLRSLLRSSPPAETLPTPVHHPPPIHPLYPYPSPSYAPPVPTPTPAPYYSPLPSYRGSPSIWAMHHHHYPLSGVRSSYPVPSYPPVTTPPWSLPYQPIDVKRDDCTSDVPLNLSKDAG

>Myzus persicae (aphid; Hemiptera)

MTRDSAAVAALDDRRRRCPISAASNVVDDASTLMTNSIAMSGSSSNDDVGGGDEGGSLKSSSEVSTTTSCCGGSSSSSRSGSDDDDSATVTRLTTAIDDSCSTTIKTPPSLHKRMEDDNNGDKESSSSSHSDEAAATTTPTNIGGRLTFYKDGKFIFQLAAHHQQNPCNNPSALTSPCRWVPVPTVIQQHKNNVKCIGEWSQHQQNKTIWPYSVTNNASESNSPQPVQQLQPPTVRIGERKQQNFQHRTTPPPLSLITGAVNTATHTGGVVVATTTNIVRRPSRKCSTTVTAENITNFHHQRCQPLTQLRGLPRRSQQHQHQAFMMVATEAIKVQRRLLSLPKKQRLWCQRKRKSMTRNSLLQELKPPQINLECFVRNLWCRRHTTELLLRRQLSINNSGDNVAATISAVVDVTPTVISRHVAVSTASVVDSTCPNTGGSKRRSMVSPSPSAAILIPITATSMNLGNKSGNSSPHKKYKSGHQQQQEQILQPQRRVVVDKIVKKAIPQPPPQVISTNDHSITAILSGGAAGAKRSGSSGAVSMVMDPENGIVTTTTTTTSAVLSPHKNLHTPSPAPLSLLRTLLKSPSSESSSPPIAASTNGVGYRHHTNGSRKRSSVESTAVSSLPSINTAAIKVENGLGTSPSAIPSVGANAADNASTVLTALHQLPTIHHPAAGQLAAAGYFNVLYHRAAMAAAAMAYQTHAQLPQPLPKSQPPSLLSSQFPTTSGASTTWQRHLAQRPPLLHPPEAIVGSGVTNTAISSSISTVVAPYSSPLIAATVNGSSLLPPATPSPPPPLMLHPHSAHRHHLLLHHQQQYQPAPTVQQQPSSFHQHNHRQQSSSLKKRSGVAIGVTEYGVCIEENNSGVVATLDEESSSSADCVPLNLSKDSIAENVSGTSPTARVR

>Frankliniella occidentalis (thrips; Thysanoptera)

MRIQEKIEASATCAKMTEEKSVNGVCEVPAAAVSKRARGYVLDGVRPVRPRSVDPDTSHAFAGGRLKFFKDGKSILELSYRRDGDRMCWVSNWPVPGATRQESSAASLSVSDDNSSVQSSPWQRDHNWKQASPRRGRGAHLQFLMRPLNAARKIRRLSFISRSIRRKRRRPYDAADMNAPLIKEEDADDSFNSSMNSTLNSSKTPSKGKLPGIVQNLLDRVKTNEPSRPIIPSGRVDPNIISPRKRILREMERVSIDESSNKRPKQRLSQGNGSASSNNSTVTCHTLHPAGAPPPVKTAASSYSITSLLAPREEETVVSKETEPSFLRNLLKSPSSQSGSDVSETNSKVKSVSKSGSVRKMSPPQQQPVTLPTLSPSPGLRSPAIPSAVGGAAHYAPYIPTHPLIYPHCLTPHPATHPAYYTGYAPPHSPGPPLWIPYMSSLQGRAAMYPGLAPTYHSPMAPSPWGAPINHQPHLDDLRKEDDMPLNLSKHAG

>Cryptotermes secundus (termite; Blattodea)

MRVQEKASTSCAKMTEERSMNGDVPSGSKKGFGHGNADGVQSPNLSSPADPDRACSIAGGRLKFFKDGKFILELSHRKEGDRNSWVPVPKKTYWPPPAAAGTPRQESSTSLSVSDDNSSVQSSPWQRDHCWKQSNPRHNIGKEMTFIMQPLRHMLYIRNLHFFSNTIRRKRRSPYDPTEVTVLDVNSTNNGKLVRLKKVGSDPAKLTTIVQTLWERVQGSSSSETVACSSSGVSTSMPVQNHARTVTVAACRMDPSIISPRKRILREMERVSLDDLANSKRHRARTTSALPHNNNNNGSSNNCIVTCQTLQPGSALPSPPQPGTKVSVSSCSYSITSLLGTGRDEEAAVAGAGIQDSEPSFLRTLLKSPSQQATSPEPSPRPKGTNKTGGSRKASPTQHRQIRSPSHHGSPTLSPSPESLRSGLRTPVVPSVGNPAHAPQLSSFLPSPLLYTPSLTPPYLPHHPSSSLGHHPYLSGLSSIPPPYRGSPSPIPSYWVNYPLSSLPRGALYPPGLVPPCQAPPLSSCPWGSLNQQPLDDFKKDDGASDVPLNLSKHAG

>Periplaneta americana (cockroach; Blattodea)

MRVQEKVSASCAKMTEERSMNGDVPSGSKRGFGLGNADGAQSPNLTATADSDRSCSIAGGRLKFFKDGKFILELSHRKDGDRTSWVPVPKKTYWPPPAAAGTPRQESSTSLSVSDDNSSVQSSPWQRDHCWKQSNPRHNISKEMTFIMQPLRHMLYIRNLHLFSNTIRRKRRCPYDPTEVTALDMNNTNNRKLVRVKKIGSNPNKLTVIVQTLWERVQGLTSSETVASSSSGVSTSMPMQNHARTVNVAASRMDPSIISPRKRILREMERVSLEDLANSKRQRARTTSALPHNNNNNGNSSNCTVTCQTLQPGSALPSPPQSGTKSSVSSCSYSITSLLGTCREDESSTAGAGIQDGEPSFLRNLLKSPSQQTTSPEPSPRPKGNKTGGSRKTSPTQHRQIRSPSHHGSPTLSPSPESLRSGLRTPVVPSVGNSSHSSHLSTFLPPPLLYAPPLTSPYLPHHPTQALGHHPYYSGLPPVPPPYRSSPSPIPSYWVNYPLSSLPRGALYPGLVPPCHPPPLSPCPWGPMTPQPLDDFKKDDGASDVPLNLSKHAG

>Clitarchus hookeri (stick insect; Phasmatodea)

MRVQEKSYASCAKMTEERSMNGGPDGPSENKRGFGLSNADGTQSPNLTASVDNDKTNAVGGGRLKFFKDGKFILELSHRKDGERMSWVPVSKKTCWPSASISTAGTPRQESSTSLSVSDDNSSVQSSPWQRDHCWKQNSPRRGVGKEMMFIMRPSKHIRHIRNLHLLSHTLRRKRRCPYDPADVLVTVDRNGTTSTRLARVKREVLSAEKFNLLIQSLWERANMASEGAGTSVLVQNHARTISATVGRMDPSIISPRKRILREMERVSLEDLGGSKRQRAHTASALPVDNSVGGNSCTTSCQTSKGSIGSYSITSLLGPGRNEEQPAAEPSFLRTLLRSPPPSPPEHSPRHRSTAPAASSAPGLRPAPVVPSMSASPHLTPFLSPSLLYPTYLPHPHTSTHPFYSVLPPVPSPFRSSPLPALWANYSLSSLPPPPRGALYPGHSAPLSPCPWGPLTPHPLDDFKKDDPSSDVPLNLSKHAG

>Timema cristinae (walking stick; Phasmatodea)

MRVQEKSSASCAKMTEERSVNGGLDGPSESNRGFALANADGSQSPNLTGTVDNDKSNSVAAGRLKFFKDGKFILELSHRKEGDRTSWAPVSVSKRTCWPPVPSTIVGTPRQESSTSLSVSDDNSSVQSSPWQRDHSWKQSVPRRGVGTEMTFYMRPSRHAMHVRNLPSLRRKRRCPHDPTEAPVSDKNQINSVSSMRVKRAGRLCKLSVLVQQLWDQNTTKIEGTSPLASTSAHNHVRTATVPSVGRLDPSIVSPRKRILREMERVSLEDLANSNKRQRATGALVTNSVPCQIPTGPLATPQPSPLQATSKCISSYSITSLLSRDEEPSPAHEGSEPSFLRTLLRSPQHTPSPEPSPRQRATPVSRSRKSSPTTSPAPGLPRAPPLVPSMGASPHPLTPFLTPSLFYTPYLPHSAPPPPYYTALPPATVSSLWAHYSLSNYSRGAGLYPGMIPTAPVGPCPWGGPIAHQPLEEFKKEETSSDVPLNLSKHAH

>Laupala kohalensis (cricket; Orthoptera)

MRVQEKLFTSCAKMTEERALNGGPEVVSSGSKKSFGPREIDGAQSPGLTPSELDRNNSLTGGRLKFFKDGKFILELSHRKEGDRTSWVTVPKKNYWPLSSAAVVAGTPRQESSTSLSVSDDNSSVQSSPWQRDHCWKQSNPQQGISKEMTFFMRPLKFVKEYRNLHFCSHNIRRKRRCPYEPIDFSFVNSISVNTTKNTKIRKSGHSRAKLSVIVQTLSNKVLGPSGECFAGSSSSSSTSSSGSLQSQARTVSISGRIDPSIISPRKRILREMERVSLDDSVSSKRRARTSSALSLNNNGSSGNCTVTCQTLQLSSVLPSQPASKATASSYSITSLLGTGQQNSLSSGTDYEPSFLRSLLRSSSQPATPPELSPKSRSGSGKGKTDSSRKASPPQAQHMPATSPTLSTSPDSHSCGIRTPIIPSVGSSPQHLNPFLGAPLLYPALTTPYLPHHPGTPSVSHHPYYSSLPPVPSYRGSPPVSSLWANYPLSSLPRGALYPGIITPCHAPPLSPWSSTSRQTIENLKPEDNPSDVPLNLSKHAG

>Stenobothrus lineatus (grasshopper; Orthoptera)

MRVQENSFASCAKMTEERSVNGGPDIPSGSKRGYSAGDIDGTQSPNSSTSEADKSCGIAGGRLKFFKDGKFILELSHRTEGDRTSWVPVPKKTYWPPPATTATSTGTPRQESSTSLSVSDDNSSVQSSPWQRDHCWKQTNPRQGISKEMTFVMQSSPHMRHLRRLHSVFQNIQKKRRCPYLPIEVSVENHSGVISVHARGGKKERSRIKLSDIVQTLLDRVKGSSSSNESSVCTSASAAIPPNHVRTVTVATARLDPSIISPRKRILREMERVSLEDMSNSKRQRARTGSALPQNNNNGSSGNCTVTCQTLQPGSALPSPPQPNVKGSVSNYSITSLLGNSRSEDVPTPGLQETDTSFLRTLLKSSTQQCSGAEQTQRPKAAPKPAGSRKSSPTQQQQQSPQQLPQTQHSQNYSPTQQQLSSPSRLSSPALSPSPDNLRSGLRNPVVPSIGSSAHLSPFLTPPLIYPQLSSPFLPHPPPTLSHHPYYSSPPPVSAYRGSPSPVPTLWVQYPLSSLPRGAVHYGLVAPCHAPPLSPCHYSPLTPQPLEELKKEDNGSDVPLNLSKHSG

>Leuctra sp. AD-2013 (stonefly; Plecoptera)

MRVQEKDSESCAKMTEEMSMNGGPDIPSGSKRGCGFRLADGTRSPNLTATSDSERTATSTLAGGRLKFFKDGKFILELSHRKEGERTSWVPVPKKTYWPPPGAAGTQRQESSTSLSVSDDNSSVQSSPWQRDHCWKQSNPRHNVNKEMAFFMRPPHHAVYVRSLGFYSKTIRRKRRCPYDIVKVNLEKEDINDVKPARSYKRTGPNPARLTLIVQTLMEKVRSVNASGDNMIGSNNLLGNHMTNCASTSRLDPSIVSPRKRILREMERVSLEDQANSKRQRARPGPSSSHATNNGSSSSCTVTCHTQVLPSPPPQLTHRGSVSNYSITSLLAPMDEAAPHSTETESPFLRTLLKSPPSPSVSPRPQSSVSIKTSPSPVPSPPESHRNHRSSHVGPSMSSAAHMSPFLTPGPSFLYPASYLPHHPYYSPLHSVPAPYRGSPHSASPLWLPYPLSSRDGIYPGLMPPCHPPPMSPCNWGPAITHHPLADMKRDDTCSDVPLNLSKHAS

>Forficula auricularia (earwig; Dermaptera)

MRVQEKASASCAKMTEEKNLNGSSKLPTGSKKPFGINSDELKNAASSEVDRSCSIAGGRLKFFKDGKFILELHHRKEGDRQSWVSVPKKTYWPPPGAAGTPRQESSTSLSVSDDNSSIQSSPWQRDHCWKQTIPRIGISKELTFFMQPLKGAKEIRKLQCFTNILRKKRRSPYDKQDVTLIKDLKYNIKNRKQIKRLKLKDLVQSLWDRITGPGSESVASGICGASSVTVQSYLRTTNTSSRLDPSIISPRKRILRELERVSLDDQTNSKRQRGRTAPASAFAHNNNNGNSNVNCTVTCQTLQPAIAVPSPVVPSPPRSTPKSNITNYSITSLLGSTRDEEQKISPHEGESSFLRTLLKSPSRPSTPEPISKNVTKLNSSRKSSPTYYHSPSPESQHSGIRTPLVPPPHLGHYLSPHLLYQSPYLPIHHNSQAFNPQNYYNGLSTIPYRGSGNTPPLWFPYPLSSLPRSPLCSGLSAPLAPYNWSPINHPPLDDFKKEDNSSDVPLNLSKHAG

>Baetis sp. AD-2013 (mayfly; Ephemeroptera)

MDDNNGIGVMRMQENIVCAKMSADDRGVNGENIACGKKSPRNGISETAATNGAFSQKPTVQTDSGGRLRFFKDGKLIFELSHQKEGERASWVPVPKKTYWTPRGKCSSLVPVATAGSSRQESSASLSVSDDNSSVQSSPWQRDHCWKQVHPRKDAGRYLTFFMRPDSRCKTFRRRNLKALKPKRRRPFDAHEVKDDCSGPPSGDSKKLIGVVRRKLVDIVEILASRAVELKKPSPTPGCSKADHSVVSPRKRILREFERVSLEDQNNKRHKRPSSTSQVKSPSPPTLGPHINAIRTKGSYSVPASTPSPTSSPSPSSSSKNKERLGSYSINSLLGRSDEPAESEPSFLRSLLKSPSRSDQGSSGSSPSCVSSPQSPSEVRWRPPKKKVASPDPPPQTHHLPYHQPFPGLGFFPPPVPPQYFSHYRGLWPPPHPSAPALNYPAGLLPLGAAWTHPHLGPSAPLQPQPYYVDAGKREELAADMPLNLSKNAG

>Ephemera danica (mayfly; Ephemeroptera)

MDRKNGIEAMRVEESITCAKMTEERSKNGVSEISSSKTKSIGSVGDTVSVAANGALSPKSTLSTEGADCQRSAGGGRLKFFKDGKLIIELSHRKEGERTSWIQVPKKTFWPPAGGGGALLSAAPVAGTSRQESSASLSVSDDNSSVQSSPWQRDHCWKQQTPRKDLGRELTFVMKPSLRMRCARHSARPAVQKKRRRPIDPIEIVWEDMENLPNSPKLRTRNIRHCEGKLNSTIERLFNLAIEASSNCSRQVVQSCTKIDPNIISPRKRILREFERVSLEDLASKRHKRPSALVKSPSPPIVPGCSNINSNNNINNNNNGSNIRPTAVKTKGNYNVPSSTPSPPSFACSNTVVSNSSGKSNKVSEKLGSYSINSLLGKCDSISDSVVPSEQSFLRSLLKSPSRSSGGCDLASVAGGSSPSSMVHSPQPCQSPPDIKWSRPHKKKVSSSPDTQSSSSSVLRLATNPSPAPPTVPSHHLSPPHFLSHPHPHHMYPGLSFLPHQPPLIPSHYYAPYPSRSPLWPPHPAAAVNLSYPHMLPTPWSHLPLGPPSSPQQPLLVEPGKREELTADMPLNLSKNAG

>Calopteryx splendens (damselfly; Odonata)

MRVQDSSTSASCAKMTEEMNKNGGPEVNTGRKKGFGPSNANGSQSPSLSIPNDTESCKTVSGGRLKFYMDGKFILELSHRKDGERTSWVPVPKKTCWPIPSSASSTTMAVPGSTPRQESSASLSVSDDNSSVQSSPWQRDHSWKQGAPRQGISKEMSFVMQPLKRSGQLKKLRHSHPVRKRRRRPHSPSDVKAQDVSDSIGLAKRASPRSKLLDLVESLWKKFGGLSDGVARTSVAIVTHNHVIKSEGVGVPTGGKLDPSLVSPRKRILREMERVSLEDHATSKRHKSRSSALVECRKPPMNTSNSTCQMQRLKAVDEVPSPPPTKGTDRLSNYSISSLLGKVSEKGSGGEGEPSPVLRSLLRSPTRGGSNSQDCHPPSPISEVAPQVMVAARGGGKVPSPRHRHASPPAVPYHHPQHSLPHHHHVPLHSPRLPSPPTLTPSSPPHSRGLSRPGMSSLSSSSSPSSSPSSHMPFLAPPTLFSGGSLFSAPSSLAPPAFLSPHPSSLAHPYYAAAAGLPSGMRQPYRSTPSPVPSLWLHNYGMPSLPVARTGDAMASRSLYPGVMVGPCHSSMGVMPPWQSLVPHLQEDYKKDDGTSDAPLNLSKNAG

>Ladona fulva (dragonfly; Odonata)

MRVQDSSTSASCAKMTEEMNKNGGPEVNTGRKKGFGPGNANGSRSPNLSTSNDTEGCKTATGGRLKFYKDGKFILELSHRKDGERMSWVPVHKKTDWPLPASASSTTMSVPGSTPRQESSASLSVSDDNSSVQSSPWQRDHSWKQGTPRQGISKEMAFIMQPFKHSGQLKKLRHSHPVRKRRRRPHSPSDVKAQDVSDAICLAKRAGPRAKLPALVDMLWRKLGGLSEGVSRTSVATLTHNHIVKPDGSAVIAGGKLDPSLVSPRKRILREMERVSLEDHATSKRHKSRSSALVECKKPPQNASSNSTCQTQRIKPVDDVPSPPPVKGSDRLSSYSITSLLAKGSSEKGEEPSPVLRSLLKSPTRGSGNSQQEHPSSPLSEASPPGMMVQARGSGGGKVPSPRHRLSSPPSAPYPHPIPHHHVPLHSPRHSSSPVAAPPSPPRPNSRGMARPGIPPMSSSPSSSASPSSHLPSPFLAPSPLFPGGGLFTPSGIPPSFLPHHPTLAHPFYAAGLTSSMGQPYRSPPSPVPPLWLPHYRMSSLPVARTGDAMAARSLYPGVVVGPCHSSLGVMPPWPSLVPQIQEDYKKDDGNSDAPLNLSKNAG

>Atelura formicaria (silverfish; Zygentoma)

MRVEDNLASASCAKMTEERSNKNGGPEIVASGKKKGLGPGLVDGTQSPHVTPSTDTDGAQHIAGGSLKFYMDGKFILELSHRKEGDRMSWVPVPKKTYWPPSTMGTPRQESSTSLSVSDDNSSVQSSPWQRDHCWKQRSPRRGVNKELSFFMRPVQNIKHVRKLYHYSHSIRKKRRQPFEPIDIIHGENNRHKNCVWVKKKDLARLDLVAVLKKLRERLSAGSDGVKMEVKQTVTVSPSAKLDPNIISPRKRILRELERVSLEDVGSSKRPKPRSSALTSTTNNSMCQTQTTVCRTSAKGDKLSNYSITSLLGKGGPSPDAGGTEPSFLRTLLKSPPRSCETGPIASTSGATPKPQARVQPWPANKKTSSCSPLHSISHLTSTSPSLSPSPDSRNVRVTVPSMGSSHLQLFSPSLYTSPINTHAGLFSPTLSSPFQPHHPPFSHPFYSNLPRGSPSLWVGYPMSSLPRGALYSSVPGSYQPVLSPWIPLASQSPDDLKKEETAPDVPLNLSKNAG

>Machilis hrabei (jumping bristletail; Archaeognatha)

MNGMRVEESPASACAKMTEESSKSGGPETTNCHKTLTEIIKVEKSGPVNKNGTRSPILTSSSEQDNPRNPPGGSLKFFMRGKIILELSHRKDGERTSWVPVTNKTYWPPSTVVCGNSRQESSASHSVSDDTSSVQSSPWQQEHRWKQSSPRLGIGKEMCFYMHPPPHVIKIRRLVPCALTNIRKHCRRPYQSIDETSFKKAEVDIKPYVFSKTKRRSKSGAIDSIINKLKSKCADRSRSEASGSKTDCSMKPPVITKASVVASVGGKMDPSIVSPRKRFLRDMEMERANAEDGTMPSTTKRHKIQVSSCSVSVGDGISAVTVSALPQSSSPSSVSRTVTSNSQSTVTSACFSSPSQLTTSCQNRSNNSGSSVSSKRSSAKNVEQLGSYSITSLLARDGIVSCEGSSKTRGDAESSPSFLRSLLKSDGLSASSSPTVNDSNKNRKTETLRSVRSPPFHSASSATSDSPPHDPRIFRSTAPSILGSVGPNIPGFLAAAAARPQFPHHYPLSLPPSLPPAFLQQSSHSSFYPFTSLLLPYALPGVAQETPVPHSFHMSPPSRAPPIGSWSPYSERMVRNVIKIDEGASPDVPLNLSKNAG

>Holacanthella duospinosa (springtail; Collembola)

MISGEEEEEGRQKSSSESPPSTATPSDEEPFLGRNGTRTPEGVLTPPIDNTPLKYNGRLEFYKEGVLCLELSHFDGGEARKWLKTKGKSQPPHVVKVSKEVPVPCSPLPVTINGAPSHDMSDEGSSLSASWSDSPKPRGKSLRKISLPLKRTSIETTFFMKASSCLPKGLFPRKPFSRHSLATATTATQIHPNPTHLCACGEETKRFIRKGTRIPRPIDVIILRLRGDSSNHLPDSRSPQKVGTPNKKRESREREKSVKKRKIGRRGFEKTILSVPQPNQDCTDGKSTPKKKESSGESLLKPIPSTAQPITSRRNSSKDKEEEILSPEEQEEIQKHAMSKAISYTIQSLLGTPPPPTTFTKLIHNSSEPKQPIPPLHLSNLFGKEHDKNSFLRHLLDTSDPIPSTNHLTNSPTHQQQNNQSRTPKNKRKGTKERKARDSDESPESQEVLSQSRTSDRGRQFQLSSDSPRLGSIGNYTQASTQDESPLDYGALLQPQIIPTPSHASFHPHHLPPTPPLSGLRFPEDNDAPIDLSQKQRKFRNTTIDSDPDVPLNLCKKRRDD

>Orchesella cincta (springtail; Collembola)

MQIMSSPSMSDAADTPVSGSGGSPSPQSPLSTTKQILDFSKKECDETDPRNTISPKLIESTESERSVNSSPATLVPTPRSTPPMNGILTPPNDAEGSSESSKSSKPGGRLKFFKDGILCLELTHHGEGERTKWTQIRTKTVFPTLVHTVKEATIPCHSTPSSFASPSSLTHNHRVESSTSHSLSDDASSMLSSPRVRESHWKQSVPQRGVSACLQLYMKPAPCALKVRRKLKDVSQDQLRKRWPYDYRRTDILLKLQNSVQCCCDVTSRVLKNNNNNKSSSSSSESSDENVTDNDTTLRKHLSKDPKSKRDNSKKISLNTVIENLEKLAESQASIKAVSKCKSVKSNGDTHELSGKRCSSTKQAAVAPKKRLFNEVEPREPSTVRPKKKSPNRSLDVGKTILNVPGLNSEFYKRHSMSKANKSPVAGPDLGGFETRHSRHMHESAVVHNKRNNRSPHFGAKSNPPGTSSSTLLKPGITTELVDHYPPVSSSGRGLSRISDEHASSIRHQDLSSQSLEQKLFQQHTVSKAINYTIQSLLGTGGDPPASLSSKQSLKEQGGYSQSKEASSHGHLQNIQKGSRHSSPINCTPTSRPDQDQKSFLRHLLDTSDPIPKATDRTNKNSSRSTPVNHNNRTSGLKTSKASSNNNHEVRTNSPGSIPKSTGRNNAPTQGSLYSIPKDPVAEAHQNLATNIALSSLMSFPMNMTPPYDLLTRGANQHSADALTAAAAALQYGSLIPGNFFPPVNSTMAAAVAASYLGVMPNLPMAVPPPTSRHSPSPKSNHNVRRNQETPNSANPRSWPGMVNGHPPGSSPPFRSTTTSTAINTNNSQNSPLRFSSAISSPVEVSSCGALNLSEKKKSHVPEPVDMDAPLDLCKKSSR

>Daphnia pulex (water flea; Diplostraca)

MSDVTMDYSGDASQADEAAQSPETCVPKSPSSPSSHSRIKSEDSSDFESKPKAPEGGRLQFYFDGKIVLELNDRRENGKTWWVPVTQKTYWPPPPPSCSTPGSTVRQESSASFSVSDESSVQSQSSPWLRETRWKNPNPTKKRTPSDVEGFVFAIKSRERCRYQWIQRRRPFRLTEQCHLGPNSECGQCKVQWKKKGLARPSVAAIAPRLLEKALSKKMMATAVPIERMESLVSPRKRLLRDMEKVRLNDKSSHSSSNVLKKAKALMPATPLPTHSAAAMAAAAAAAMAHSLQPSKVYDRQSSYSIDSLLNKERQEEAAAASCSSSFLRSLLRKAPQPAPNVPTVVNGVRNKSLVSHQRPRDVTVSSVGGSRPLQPPVWLPYPTPFPPPPNSMTPGSSATLAARPGPSSSSRRETLGDRSSRVPTPPDYDGDDDVPLNLTIRRAREHS

>Eulimnadia texana (shrimp; Diplostraca)

MMSETTPSGVLLVQDPQQTDSRSPASDTENVPSETAIKTEEDCKTNNCTKSDSGGRLSFFLDGKLVLELSSVSDGTTKTPWIPVSRKTYWPPPCGSSFSNRHESSASYSLSDDSSLQSSPWQRETRWKQPLPRPNASTALSFSFLASRPRSVFCTCRSRRRPWLLCTCHSINNVKSGYKTLSESKPRPVIDSILPRLTEKARSKWLGSPLSRKREKQTKVYDRQSSYSIDSLLGRERPETSASPSFLLSLLSKTSNGMHSPRLPASRAGVVPFFSSGQAFPASGVPHPLWMPFPLSHLAQLPHNQAGLNSRQAAPISQPPTPDTEEEGPLNLCTSRRERDDDS

>Triops cancriformis (shrimp; Notostraca)

MKCCEPPHSQLIMSEEQDIEAGLMKPKIEEPLLVQSTAPIVSPTLAKHSEVENNGSCSPFTPEPVAEPKPNGSRLKFFFDGKMVLELSDKKDGEKVSWVPVTKKTYWPPPLPGRPLRQESTTSYSVSDDSSVQSSPWQRDTRWKQPAPKRARCQDRTFYMLVPKSERSRRCSSHVQYRKPFLVQLPHCQCKRTTTRPRGAEPTKRRLSLPRMVQKLGERASAMSRAAEFSTHSVRIKEEWRPCEIRSAKDFHLTAVLGSPEQSTRKRAWKESDRARSADAYLKKTKAQQPVSALVAAPLALSSNSKGFDKHSSYSIDSLLAKDSLNSASANPEVNHGPSFLRTLLKAPASSVSPASGSSPSSAAVTLANKVKVNSASPSRNGHSKGDSASTPRHSPRSQVPTPPQDAAGLQLPRTSGGGTPRPSYSASPGPRFNPAVLTGLLPRMAGIPGLPPPWEMYNPYMTQLLGASPSWMSYSWPPPLGALPSAAFGFSEGLMMMPPNAAPSAIGHKTNGASIAESDNCSSSSSSSNSRSVIVKPSSTSKESRKSVEDALIQDVPLNLSKSRSGESR

>Caridina multidentata (shrimp; Decapoda)

MKNKTETLIGNEMSKPALEGGGGGGGLGVKMEKQPPSPVRSPPPSPPHLGVRPSSTPPSSQTPKIKCESPVSIGSSHASSPRPVGSPPSLGPGAEVKGKPGTPIPSGSVKGVAPRPGSAGCGEDASKGACGTSAPGGGRLTFFKEGRFVLELSHRTEMGAPPGWVAVKSKTYWPPPSSATTTTIQHPLRHDTPTSQSDDCSSLNSSPWTNEHSRKQSVPRRNKASVQSIFCWQAKVVSGDLNKRRRAFRRNPFIYPAGCLEPAKQKSQAKTVKSESCTDVTDVKKANLESKVQSLEDKLGMKVDRKVSVKVLFEKPPASTSVEPIDPTFVSPRKRYLRLMEDQDSIHRKKLHAVANSTPQIPGERPSSSSSPAHSFSNRSAINSQTSSAYSIDSILNSESANRKNDSFLRTLLKPEPKPGVPQSKSVIKERVNNEVIANTVKVERPERTDRGPPDRLEQLSTPYKDRKDPVADLNRLAAERYNLQGYGSLGGSLASLYGLNLLDPRYMMLPSLAPPMSSQQTEMAVAAATAAAAAALSTYPLSHLHPSLAGTQYSHLLSAGLPSGYPPAAFGQLLRPQPQPSRLSPHPLAHSPSPKPPQAPSTPASPHMSRGASPRGASSPWQPPQPAHIFSPQKPIASPPSPPPQDAPLNLSKPRHHVEK

>Homarus americanus (lobster; Decapoda)

MKNKTEVSSTDVCVRMSKAGLEGGGLGIMEKRPPSPHSPPLSPPTSATTPPSQTKIKCESPVPLLSSHTPSPRPLVSPPNTGPGAEVKGKPGTPIPSGSTKTVLPRPGSASGDEAKVSVTTGSSPTTTGGRLTFFKEGKFLLELSHRTDMGAPAGWVPVKSKTYWPPPSSATTTTTQHPLRHDTPTSQSVSDDCSSLNSSPWTGEHLRKQLTPRRNKAFVLQQVSFHWHALPLTRDIRKRVRLRRNPFTYINGPVGCWNPSENAQVSRRTIKSESYESALEERKHKLETRILVLVSKLGLKVDPETSIQALFEKPVASAEAVDPTFVSPRKRYLRQMESDQESIHRKKLHSGANSVSHVPGENLSSSSSSTQPRPPPNRSAINSLTSSVYSIESILNNETANRKNDSFLRTLLKPEPKPSGTQSKSSVVRDRVNNEVLASTFKVERQERTDRGAADRLEQLSAGQIERTDSLVDLNRLSTERYDFSRYGPMASLYSLHPYLDPRYMMLQSLAPGLVSPISSQHTEMAQAVAAATAAAAAAAAAGLGTYPLGHFHPSLTGSQYPHLSGSLSSGYTPPTSIAAQLLRPQAQSSRSSPHAMSHSPSPKPHIPNTPASPHLSRGASPRGASSPWQPHPSVHPHSPHRPLSSPLPSPPPQDAPLNLSKPRNHMEK

>Proasellus hercegovinensis (isopod; Isopoda)

MNISKNLNVSEGSKGTAKISENDDIPMFLSKSPPRVNALNFSIKREKLFADEESSLNDGKLSPPLSSLGPGAEIKGKICHPISSSRNDNIESPEDGCKVVPAGVSGGRLTFFREGKLLLELSHRNDQGGAWVQVKSKTYWPPSSTATTTSTQHPLRHDTPTTHSVSDDCSSMNSSPWFNDHIRKQVAPKSTNKIAFTSGELFCIQRSQYIRAIRKKVRLRRNPYIKIKVKTSLDINFLKDSSKTFQSHCHDMKFSDKRKRLLSFIEDLEEKCSLKIDNNISIQTLFEKVDMVNNDCVDPTFVSPRKRYLRSLESEIDSNIRKKVHSSYNSSVTNSNYKHCDRLSSSSPSYHTPSKSAINSQNSCSPYSIDSLLSNDTENKKSDSFLRTLLKPDSKVRSPIPKANLIKERVEGVSNHIKYERGSSSSHERDRKEFDYKNATMADHLDMSTDLSRFPSPNLSGLYLNPYFDPRYMMLHSLSGAPPFIPAVAQPSEMAVAVAAAAAGYPFNPLHISALASGTLMSPGSLCSTYPPATSIASQLIRSPTPTRSSPLSVTHSPSPKSSQVPSSPHLPLNPHLGSMSSSLRDVVWSQPSHNNSSHKQSSNSIPQPPQDVPLNLSKPKGKS

>Hyalella azteca (amphipod; Amphipoda)

MSQPAVGVLAEELVDCQKTSFANSIEAEPCSPTSRSFSPAISDTEICERREEGTAVVAALPPASPIKHEVLDLTSSNRTPFMDLSVKPKLANQVPPVSSMLPSALLGSTRSVSNPTEVLTSTSSFHSRNNSSKCVISVNASNVLGNPTCASPNDDHLRNSVDLAAGLITIDTSSLSSIDAASLSTSFHSKSAVQKSNKVTAQGTIVPPLANSIQDSAILQDSSSSSSASAVLSSSIPSSRNQEISATSQDHLGQADQFCSSQIYSELPFSGSEKKNTLKLNCASKLLTSHSKQVSEPCSDVFTNGHEHHGTSSTSFPASASSSPCRASLSPSPSCSESQGIASGAGSEVRGMPAIPVPAAPGRPKEAQDPSASGSMDNTDGGRLTFFKEGKLLLELSHRSANESCAINKDGKKTSGCWVQIETNTFWPPSSCRANISTPLHFPFSGYGKAARRKIRTPRRCEQQSNGVVQPPLSCSASPTKDDASHRCVGAPVPETFLFVCPELVRAFRRSYGPLIARRPFHRFSEMDALLELDDTRKQALLSLMRECRNLLRRSSSQEGNRILQKHKLEKYSKVLAEQQNLKIDSTIKVGQIFERVMLKTPSPNNFVSPRKRYLKQFEADHEIQNKKKIVMATSSASRPASYMTDSYQNYTSSKNVSARHSALNNYGAAHTIESILNHSEKRNESYLKSLFKSEGKKSPRSSSRSPLTLESVKQEPEMLMSVKPDRDASRNSEMRSSRDCDERQSLPSYSASPHSRSSPPSPRSARASHASNTSVKNRDQPDLPYPGYGMTPQTLASLYPQLYDPNYFFINGPLAAGGLMASPHSQMAVAAAAAAAAAASISSMSSYGLAPLQLQLAATQYSSLFNPVAFAPAIAAAAAASPLSPSIAAPLTPALTPGAVALSMPHMNSSTLPSAHNPSLSLPHVVTSHHPSFINSVASHQSRSSPYPTPSPHPRSPYSPSVKSFYPPTRSPQSRSRSPPSRSPVMMSTTPTSYHPRKPHPPPHHYPQPAMHPSSPWRPVAAHQHPPPTLSSSQDTPLNLTKPRCQ

>Oratosquilla oratoria (shrimp; Stomatopoda)

MKNQSQVAAAAATNVCVRMSKAGLDGALAIMTEGVKQRAASPPPSPPNASRSPLVASATPPITIRIKAESPTLSLDSKAPSPPMSGPPTPTPSLSSGSTHVSMGPGAEVRGMQAKPISSDSKPGTPQSGSGSGGVEVEGSRGGPGGGRLTFFRDGKLLLELSHRSEGGTGGWIPVKSKTYWPPPSSTPHHPLRHDTPTTHSDDTSSVNSSPWAVDHIRKQSITQRPNSVLVELLSFSVCVSPSVRGLRRGVTRRRNPYKSIKGPMGPLDSLVQIERTKENHLSNKSAQERRSLLEGRIQMLGERRGVKIETSVTIHGLFEKPIVTLVENVDPNFISPRKRYLRQMETDSEHSLLRKRVHAAASNSVSVSSHLAEYPVASPSPSSSSSSATSHKSAISSQASSVYSIDSILKGETANKKADSFLRTLLKPEPKSSVTPKPSIVKDGVNPMSPVLKTEIIDRPERMDRPELNSASKDRNDTAHDLNKLAAERLDLSRYGTPSSLANSYLLPYLADPRLMMQSLAANSGFVTPVTAPTTEMTAAAMANVLGPYPFQAIAANPYHHLLHSALPVPAYGNPNQALVHRLLRPPTPNRTSPYPPSPKPAVSSHGPSGQVPPSATSPHMPHVPSPHRMSPWSLSSAHSPHKAAQSPVPSPPPQDAPLNLSKPRSSSGK

>Euphausia superba (krill; Euphausiacea)

MKNKTEVPSLDVCARMSKGGLVEGGGVLEKPKPLSPNSPPQSPLRGLTPPTVTQLKIKSESPVLSPRPIPSPPSSEPGAEVKGKVCTPIPSGSVKLDTSSPRPGSADDSKANINVSGNSPGGRLTFFKEGKCVLELSHRNELGGPGGGWIPVKSKTYWPPPSSAASVTTQHPLRHDTPTSHSVSDDCSSLNSSPWAGPDHIRKQSAPCHSKTKIGLAQFFIKVSSVSRTIRQNLRRGRDPFRRIEGPKDIKDISKENIDQSKENIDQKDKNNEQNILVLKHSKELGKAKLENKVNSLCERIGIKMEVGISIQTLFEKPPVIVPSVVEGVDPTFVSPRKRYLRQMECGENVENFQRKKHQTGAGANNTGHIVDRASSGSPHAQLHHSISSPHRSAIPTQSSSAYSIDSILNNESASRKNESFLRTLLKPEHKSSGISLSHKPSALKDPSSLPYHSQHQSQSRNSNSSSSGGSSSGNSSSGASDLLGIGIKSERGSVEHRSELNSNSSRSGDKDRKDTIAELTSKFAQERDYLARYGSMSGLASLYAADPRYMMLAADPRYMMLQNLATGLLPPIPAQPSDMASAVAAAAVAAAGAAAGLGSYPLPPSLMGAQYPQHLLAGYNPQTSLAAQLLQQHQQQQQQQQQQQQQQQQQQQPHNSARSSPHPRSSPHPRSSPHPMPHSPSPKPSAAALSAPSSPVHHLSRGGASPRGASSPWHTPPPAHPHSPHKHPSMSPASSAPSPPPQDAPLNLSKPRSQLEK

>Lepas anatifera 2 (barnacle; Pedunculata)

MTARQTVEPELRPPAPLSQLLPPATLLSPAPSAAASPAPPGCVTPPLSEPDQRAETAASGGTLKFF**G**KDGHVLLELSHAPVSYERRTSWMGVSRKTYWPAVTVAAPVDDIAGRCGSARLGESSPQRTLRRRRKQAAPLRRQALHPPLLQSGCLPAGRAQLRRLGRRPTADPAAAGRLGSGAARRRRCVEPVLSAEARSACLMSVVRRLWDRHPRTQPPAPAAACARTEKGPSRDGAVRPLSPRKRPLSAREMRVSPRPRPSPAEAPPVSELQSAAAAPTTSARRVSSVHSIDSLLSRTAAPARPPESTRTPPVVPASPVRPAEPPLTAGFWPLGYGAPYSPYALYGRYSGVGGLAPYLRAGEWLGEASVAGASLVPQPSGSVGLRRGRVRRPTPLLPAQLAAAQPSPAHRAPVHRPSHREAPVGPAEGADSWRYCAQNDTPLNLSKR

>Acartia tonsa (copepod; Calanoida)

MSTQLSASLPGGCGVVIPPSSPYRLTAAAATAGGASPPHSGGEQKLVNRQLTAVVTTISSSFTHSSHSHLENSIMVKTERTIDDDGVDLSYDSATSAVKKEITHPGSSHPLDVIHRLKSEGLNMGTGSAHSTNTTSSHFYSQPQHSPQSAAAAGSHSMPQQPASPASSRGSVGTNAGGSGGTSSTPGGRLKFFREGRMLLELTHRGVGEGERTHWVPSGNKKVYWPPGPAPLTSRLDSNASTNTVTGSDGSLETGSSHNSLTPGHPASPWGGSPSPRLPSTWQPTPTPADKPRIKPIPTKITKQPSSDIVFAMKPPQGVKNYRKKIRPFRRRCRQPGTDIKIGTFVLKLEIQSAAGKVKLKRKSPPHKTIEQRALRLDEQIAKIVTRKSQIQVPISAWRVGRTDLAGLIKNPPSEPGSPVGKMAPASPAAPSMHYPAVRRPVSPPRIYPSSPVPKYHGVTQLPGGGMKVKSELKMVTETCTSTITSVSTSQPRQNKVKAETRTVSAETRPVPVRTMPITPTIGRVKTEGGGMGHSPAVGHPGLEGVNNSPGRGLKMAALTSLMTPEMARVRMEIGPTPHHQDREPSFASPRKRFLKDYPDGQSPAKHRRLSTEHRLSTDSLGSTGDRASPYLPHRGNSPLPHSAASRTAAGGSNSQDVKPFTAFSIDNLISGSQAAAAARLAASSAAQSSSSPSRNRRDSNQYLGSPHPGLGQRSTTPSNPVGGLSLRHDLTPSSHHHSVPDSPGKVHAPTPTRPRGTSRDSLPPARPASSPPHGVSSSHHSAAASHQNAAGALMNPFLYPGGGPGAAGQHPLLAAAASQQIAYLAAIQNQANLLTQQLREQGVNPAMLQHALQQQQIQQQQAALAAAAAGGHSLFPGMPPSAASQFPGANPFAMHPAFNPYLAAVSAAGGFAGLPGAASMPPAVSIASASIPSIPTSVWTPPVGGVATPTPPNSNSSGQRETPTPTRTSHSGLSSPGYPASTSQSLATSMGGGGHHPSLLRIKESMSPPPTSGAGSMDIKREGSQRQSSEAPLNLSMRPTTL

>Eurytemora affinis (copepod; Calanoida)

MSTQISQHSGGGQTLQPRPLGGGGPPPPVSSPFGGPPIPSSSPFNQKQLIGLSSDKMVVKTERTLVKTERNVDDGVDLSFDTSVKKENFEIKMELNVETLVKTEQENEMNSSPRTPNGGRLKFFKEGRVLLELTHRCVSDGEKVSWVPTGSKKVYWPPGPTPLPGPAPLTSRLDSNASNHTVTGSDGSIESGSYSSIPPHSPWGTPSPRPVSWQPSPPHRVKSTPVRLCDPHQDLLFAMRPPQGVRNYRKKIRTFRRKCRQPESDIKIGMFILKMECQSAVPKPKKRVVVPQPASARNARLETVIAKIITRKSLVHVPISAWRSGRTDLQNLVKNPPSNPGSPSGPRSSPLPKGYPAHPGPKYPSSPGMRIKSELKMVTETCTSTLTSISSPPRHSAKVKAETRTVTAETRSIPVRTLPITPTVARVKSEPDGGHRLKASAITSLMSPELARIRMEIGPAQHERDQSFVSPRKRYLREFENGGGSPAKNRRLSTEHRLSTDSLGSTGDRSSPYRGPGSPQPPSRPGTKAPFSSFSIDSLISGSQAPSSPSRLVSRRESGTFHPVPRSQPTPGSPSRGHMLATPTPNRPSPTPARTPPPQHLGYPFMYPHHPLLAQQALLDPRLAHLAALSQASPAHLAALSASPAHLAALSGGSPFTGGVPYPFNLYQSFNPYLAASTAYTPPTPAPTVWTPPVATPTPPNSISSQRETPTPTRTLTSPNYPPPITSMPQTSVYAPTASIPPHPVSAPRPVPVNTEKIQDAHKQPETLHFQSVQSHSVDDAPLNLSMKPTSL

>Oithona nana (copepod; Cyclopoida)

MSVSIKAEATIPISEGCLKKMINGSSSSVSPTGGGGNLTPPSPLEATVKAETSRIMSPNLTKNTSSSQAASNPNSDVSESSSSGQKRIQNNNTSSYTSGGRLKFYKEGRCLLELTHKSNQEGSENWVPIAKKVYWPPPGPVKAESSPSVQSASEAGSDVGSSSPSPWQPGVPSTAPAVAPLTAQSLIRPSRKNRGIKPVAFRKQPLNILQELCFAMKPPACIVSLRKKVRKLRRKRRRPFDPILVGAFLMRIEAQAVKTRPRRKPFEMDRIKQASRLERTVTKLGATAKKAPPPKLVQIGTWKPKNSSSSSSMSQTSSSKNGSIGLNKIKSELCPVTETCTAATLPPATPSTPNHKHRVKAEKTVSLSTASGLMAATPTAAKTLPITPTMARIKRDGGSLSAGGHSIHSSEFQSSSTSLISPDMQRVKVEIGHSPFNFRNTPPAEMPRLPTSPVSQMDMSGSPHFISPRKRSLLRDYENGSPSIKRHRMSTDSRGSSSEGGGAGSPSLGASPPRHNNGGRVSSFSIDSIMSSSSSSTTNNASGGGATVHRPVPLPASPARVASKSPPVPSTPLNIPITPYSARGTPQAPGGGAMQLNPYLAHLAGVAANPSLGLSPSTHAQAVINYQYAALLGAQMYAQHAINATPPTSQHSRSSSSRQSSGNVSSPHGERPFSPLGVPQYRGEPKVATPTPKATPMDIPSRLVPTESGELPLNLSVKK

>Tigriopus californicus (copepod; Harpacticoida)

MPILSGQGSDRQQHDDSFKVMSSSTATSRDDGGGGGQGGGGNGLAKSGPQSPPNPRQLLSRLAQAEAAVRLQQQQQQTPPKLSSLGEPSSPFPLSQLASSPLGSPGRDHILLNGKRVSSSMTTMSSCSATSALAAAISSTKTSSITSQDGVLVGAHLKTVSTKISSSISSHVVVKEEEDRGSPSSLVSPSLTIDERSDKENEEDRQKGSSRQDGDNNKKVQNNNTSSYAPGGGRLKFYKDGRCLLELSHRSNGDQSEMWAPVTKKVFWPPTSASVGGSNPPSSLPPTTTTPIPSSTLVKSENGSALSDTTSTVDAASIPSPWHTSAAMLHRAKSDPLRLDLAKLPGDVRRDLSFVMKPPPAVHSIMRKFKHLRKKHRRPYQPIVIGNLLMKMETQAVISRVKRRPYEMDRIKQASRLESFVSKINSLPKKKPIFQIQSWDGTKLDNNPRPAHPPSKRMNTNTHVGNGPNLISTHPTTGVTRVKSELCPVTETCTAATMPLPSQTGGGSRNSGPNKMKAEQHRGSLGKVPHMSPYLPGSQKMMPITAIPARIKSEVGPQSIISSSHVMPDMRRIKMEISGNPGSGLLNRSLPNMSVSHSMSPGLDPREPSFISPRKRLLHHSFDVGKNGSSPAKKHRLSTDSRGSAGSSDGTSSPHVPIRNACQSPRASSGSPFNLPALAASPPRAPVGKSNSFSIDSIMNKGDQENRLVRPRPIPASPARVSDIKRDMDMESNSSRSPTPASSVMSGMRASPVRTRSPPNVQLSTYAARSGAEVPYIDPYVAHLAGVAADPRLGLNHANSNLRVPYSLGSMMNPLFHQQMLANNAVSSLAASQLAAVSGLWTPPVVTSSVANLGMGNPTSQASPLPSHLISTPKTMHGSMMGGDRVLGSGNSWSTPQYRGAPRVATPERPIDVSSTDDVPLDLCTKR

>Lepeophtheirus salmonis (salmon louse; Siphonostomatoida)

MMTKEEPASSLEEDSTAGRLKFFKEGRCLLELSHRNGRNWLPITKKVYWPPSLPYSEVKREILPSCPPQQHEPHVKSSPCVSKAPRKCSHPQRCREGAYPGLFVMKPPPQVISLRKKVRSLKRKRRSPYSAILVGAFLMRVEALSGPSVYVPRKKPQQDRIRKSSRLERLISKLPVTRISIKSWAPPSLKTVKSELFPISESTSSSSTGTKLPTKVKASTKKIHADTPRPSKTLPVTPTLARIKSEPSNHLHSSGSLLTPDMKRVKVEISGPSSPDRSHFTSPRKRALQQESLAETSPSHKRHRLSTDSLGSRDGSNSPYPNNNNNVSASPPRHHYNHHHNGRMSSFSIDSIISRDEKEARLIRPTALPASPSRLGDLRREFSIEHDGLSHGSRTPTPASFHGSPAPSRRPLSPPSTSHTSTPLVSITPYASRNVDHSHMDPRLGPQLAHLAGLAADPRLGLTPTGGPYSLVNPYSYLLPLMQGTAAGSSFPGTGRINWPPSSASVNASHRSSTSTAVAWPGQYRGEPKITSMEVGRQSPEAPLNLSLK

>Argulus siamensis (Arguloida)

MQLNCIMSQNHQSSNSSVIKVEPCEAILNGDTSQSSHSSDDIPKKDKSTSDNLLYPGTSVVDRGRLKFFFGGKIILELSHQPEGDNRKWVQVTDKTYWPPLKHVNPKRQGSSASQSVSDDGSSVQSSPWQRDHRWKQVNPKKHLSYGIDFMMHTNKHISLIRYKLLAAITKHKLRRSPFLKMEVLSHHNIVNNYQKISYRKISTCRKNIGSYIDILRKKVTNNVIAPSKDSSEHCSKSVTEVKHKLHSMISPRKRILREIEKDELQSKQIVSCENTDMNYLVSQCKKEFSKARDYSIDSLLNRESVSSPFLRSILAADETASKSDMGYVYLPHSESVMFNVPQMHYNNAMDESIPKSSVLSSGRCNQSVMNPSENILSHKANIDFQQVAPIVFNFSKSPSQLFPRSQVSNVNPKIALGSNHTLPFYPIPVQDAPLNLSLHPKNYQKEG
