## Supplementary file 3 for "Hairless as a novel component of the Notch signaling pathway"

**Supplementary file 3.** **Full-length Su(H) sequences**

>Ecdyonurus insignis (mayfly; Arthropoda; Ephemeroptera)

MQEDGGSGQAMTDLSAAGPGDQQQQQQQQQQQQQQQQQTSAGASGGGGGGGSTSHLTNGHAQSGPGPYPDNHPVDLSSPRPHDALDNGTTYHRDRRSADQYKQNGVDSEWQSPEVKSYGYPRWTVPWPPHASHSPRGTYWSDSVEPPNSSDPGAPMIPGSLTPPDKMNGEHPHHGAPGAPHPGAAAAAAAAAAMSHHFGLGAASMAAHGLQTPPSPIPTPSPPVPIERFSSLLYRPKDQRLTREAMKKYLRERSDMVIVILHAKVAQKSYGNEKRFFCPPPCIYLYGEGWRMKQEQMLKDGESEQGAQLCAFIGIGNSDQDMQQLDLNGKQYCAAKTLYISDSDKRKHFMLSVKMFYGNGHDIGVFHSKRIKVISKPSKKKQSLKNADLCIASGTKVALFNRLRSQTVSTRYLHVENGNFHASSTQWGAFTIHLLDDNESESEEFTVRDGYIHYGSTVKLVCSVTGMALPRLIIRKVDKQMALLDADDPVSQLHKCAFYMKDTERMYLCLSQERIIQFQATPCPKEPNKEMINDGASWTIISTDKAEYQFFEGMGPVRAPVTPVPVVSTLHLNGGGDVAMLELTGENFTPTLKVWFGDVEAETMYRCQESMLCVVPEISAFRGEWLWVRQPTQVPVSLVRNDGIIYATGLTFTYTPEPGPRPHCPAADDIMRNVTSHVNRVSGGPLGISYEQSASAQGGM

>Calopteryx splendens (damselfly; Arthropoda; Odonata)

MCGKRSPNMAMPSISPRGWNCRVSGCLGGASSPFRSLPQDDGDQGAVISDSAGGAHCEEEAGGGAVEEVEEGRRLEQLEVGGGRGPPGSGPYTDNHPVDLSSPRPPDAVAGDGAAYQRPRGLPENNYKANGLLADAEWQTQEAKHYGYPRWAVPWPLHSSHSPRTASYWPDMAASDASLAAADPSLHHMIPGSLTPPDKMNGEHPAMHQHHVPPPPAPPGSLHAPHHVALAAGASGGRGPGGAVSGPLLGPPSMAAHSMQTPPSPIPTPSPPVPMERFGSVLYRPKEQRLTREAMKKYLRERNDMVIVILHAKVAQKSYGNEKRFFCPPPCIYLYGEGWRLKQEQMLKDGESEQGAQLCAFIGIGNSDQDMQQLDLNGKQYCAAKTLYISDSDKRKHFMLSVKMFYGSGHDIGVFHSKRIKVISKPSKKKQSLKNADLCIASGTKVALFNRLRSQTVSTRYLHVENGNFHASSTQWGAFTIHLLDDNESESEEFTVRDGYIHYGSTVKLVCSVTGMALPRLIIRKVDKQMALLDADDPVSQLHKCAFYMKDTERMYLCLSQERIIQFQATPCPKEPNKEMINDGASWTIISTDKAEYQFFEGMGPVRSSVTPVPIVHSLHLNGGGDVAMLELTGENFTPNLKVWFGDVEAETMYRCQESMLCVVPEISAFRGEWLWVRQPTQVPVSLVRNDGIIYATGLTFTYTPEPGPRPHCPSADEIMRSGQTHLSRLPAITDVPGIQYDQQPSSQGGM

>Atelura formicaria (silverfish; Arthropoda; Zygentoma)

MTDLSTAGPGPGAPGSQQQQGSLHLNGHVGPYPDNHPVDLSSPRPQDALDAGYRDRRLPDQYKQNGLVGVPEPEWQTPEAKPYGYPRYPGPHMIPGSLTPPDKMNGEHPHHSHSHGPHPGSAAMSHHFGLPSMAHSMQTPPSPIPTPSPPVPIERFSSLLYRPKDQRLTREAMKKYLRERSDMVIVILHAKVAQKSYGNEKRFFCPPPCIYLFGEGWRLKQEQMLKDGETEQGAQLCAFIGIGNSDQDMQQLDLNGKQYCAAKTLYISDSDKRKHFMLSVKMFYGNGHDIGVFHSKRIKVISKPSKKKQSLKNADLCIASGTKVALFNRLRSQTVSTRYLHVENGNFHASSTQWGAFTIHLLDDNESESEEFTVRDGYIHYGSTVKLVCSVTGMALPRLIIRKVDKQMALLDADDPVSQLHKCAFYMKDTERMYLCLSQERIIQFQATPCPKEPNKEMINDGASWTIISTDKAEYQFFEGMGPVRSPVTPVPVVHSLHLNGGGDVAMLELTGENFTPNLKVWFGDVEAETMYRCQESMLCVVPDISAFRGEWLWVRQPTQVPVSLVRNDGIIYATGLTFTYTPEPGPRPHCPSADEIMRSGQTHLSRMPAITDVAGVQYDQQPPPQGGM

>Machilis hrabei (bristletail; Arthropoda; Archaeognatha)

MTDLSTAGPGPGSQQQGTLHLNGHVGAGPGGPYPDNHPVDLSSPRPHDNLDVAYRERRDQYKHNGLGVVGIPEPEWQTPEAKPYGYSRYPGPHMLPGSLTPPDKMNGEHPHHGHGHSHSVHSGGTAMAHHFGLATGAALGISSMAHSMQTPPSPIPTPSPPVPVERYSSSLYRPKDQRLTREAMKKYLRERGDMVIMILHAKVAQKSYGNEKRFFCPPPCIYLYGEGWRRKQEQMLKDGETEQGAQLCAFIGIGNSDQDMQQLDLNGKQYCAAKTLYISDSDKRKHFMLSVKMFYGNGHDIGVFHSKRIKVISKPSKKKQSLKNADLCIASGTKVALFNRLRSQTVSTRYLHVENGNFHASSTQWGAFTIHLLDDNESESEEFTVRDGYIHYGSTIKLVCSVTGMALPRLVIRKVDKQMALLDADDPVSQLHKCAFYMKDTERMYLCLSQERIIQFQATPCPKEPNKEMINDGASWTIISTDKAEYQFFEGMGPVRLPVTPVPVVHSLHLNGGGDVAMLELTGENFTPNLKVWFGDVEAETMYRCQESMLCVVPDISAFRGEWLWVRQPTQVPVSLVRNDGIIYATGLTFTYTPEPGPRPHCPSADDIMRTGQAHLNRMPPIADVTGVAYDQAPQPQGPL

>Catajapyx aquilonaris (forcepstail; Arthropoda; Diplura)

MTELSPVGPGPGGQQQQQATTQQQQQQQPPPPQQQHGLNGHVYGESPTPTSTANNPVDLSSPRGSELEEAYQRAERREHAYRHHNGLSVGVGGTLQDLDPHSAPMIPGSLTPPDKVNGEHHSHHPHHTHSHPGAAPPLPHHFALPPSGLGAMGPPMQTPPSPIPTPSPPVPIDRFGTSLYRTKEQRLTRDAMKRYLRERGDLVLVILHAKVAQKSYGNEKRFFCPPPCIYLFGDGWRRKREQLLKEGEAEQGAQLCAFIGIGNSDQDMQQLDLNGKQYCAAKTLYISDSDKRKHFMLSVKMFYGNGHDIGVFHSKRIKVISKPSKKKQSLKNADLCIASGTKVALFNRLRSQTVSTRYLHVENGNFHASSTQWGAFTIHLLDDNESESEEFTVRDGYIHYGATVKLVCSVTGMALPRLIIRKVDKQMALLDADDPVSQLHKCAFYMKDTERMYLCLSQERIIQFQATPCPKEPNKEMINDGASWTIISTDKAEYQFYEGMGPVRSPVTPVPVVHSLHLNGGGDVAMLELTGENFSPSLKVWFGDVEAETMFRCQESMLCVVPDISAFRGEWLWVRQPTQVPVSLVRNDGIIYATGLTFTYTPEPGPRPHCPSVDDIIRTGGVGSGVAGPVGHLSRMQSLADASAVVQYEQPHHPPQQGAM

>Holacanthella duospinosa (springtail; Arthropoda; Collembola)

MPQIMDPTDRVTSPWNQFHVGLGNSYVGSEQTGSNNGTPADSWDVSPLQDEYKFTPFNNNTTTDSPSTPASVNQYYLQQNEICVESSDQQQSLSELSSNGQQQEQQELNLNGHIDYGQSEVGHGSGLIPVPVVSGSIVVPNSNPVDLSNPSPSRHPVLDERTQQRLHHNIHQTQHDHYKHLQPLGAILHNLDQPHTSNFVPGSLSPPERMNGNDPSLLHTHPHHLSHMSGSPALSTHFGLPHAGLMPHPVHTPPSPIPTPSPPVPFDRFSSSLYRGKEQRLTREAMKRYLRDRGDMILVIQHAKVAQKSYGNEKRFFCPPPCIYLYNDGWRRKRDQLLKEGETEQGAQLCAFIGIGSSEQDMQQLDISGKQYCAAKTLFISDSDKRKHFMLSVKMFYGNGQNIGVFQSKRIKVISKPSKKKQSLKNADLCIASGTKVALFNRLRSQTVSTRYLHVENGNFHASSTQWGAFTIHLVDDNESESEEFTVRDGYIHYGSTVKLVCSVTGMALPRLIIRKVDKQLALLDADDPVSQLHKCAFFMKDTDRMYLCLSQERIIQFQATSCPKEPNKEMINDGASWTIISTDKAEYQYFEGMGPVRSPVTPVPVVHNLNVNGGGDVAMLELTGESFSPNLRVWFGDVETETMFRCQESMLCVVPNISAFRDRSDWLWVRQPTQVPVSLVRSDGIIYGTGLTFTYTPEPGPHRHPCQPPDESMLRNSLAPPTLPHHHHIPLPVPQYGPIMSSSSSGRQPPSHMPMLHDGSSSTGQLHYDLSQAAAL

>Hyalella azteca (crustacean; Arthropoda; Amphipoda)

MTGLSGLPALSTPVSSYQDLPHQQLHHSNTPNPPPHHQHLDPQHHSAPNSQHNPPSSHHQSLGHRPVDLSQAPSPRPTHQPHHPHYHHLTAPAPQLQHHLHQQQSHTLPKSEPGCAAAMLAGSLTPPDKLNSDPQQQQQQQQQHPALTQHQTIPQHPQHPAQHHPSVSGPPSLPHHHPHLLPPVMPALHTPPSPLPTPSPPHYERYTTMKEQRLTREAMQRYLNERGDQTLVILHAKVAQKSYGNEKRFFCPPPCIYLFGDGWQRKRHEIARTASSEHDAQLCAFIGIGNSDQDMQQLDLNGKHYCAAKTLFISDSDKRKHFLLSVKMFWGSGTDIGVFHSRRIKVISKPSKKKQSLKNAELCIASGTKVALFNRLRSQTVSTRYLHVENGNFHASSTQWGAFTIHLLDENESESEEFTVKDGYIHYGATVKLVCSVTSMALPRLVIRKVDKQMALLDADDPVSQLHKCAFYMKDTERMYLCLSQERIIQFQATPCPKESNREMINDGASWTIISTDKAEYLFYEGMGPVRAPITPVPVVNTLHLNGGGDVAMLELTGESFTPMLRVWFGDVEAETMYRCQESMLCVVPDISAFRRGWQWVRQPTLVPITLVRNDGIIYATNLTFTYTPEPGPRQHCPQADAIMRPHRAPLAAAQQHGDLPGLQSSPPANPLVPLQFEPSQPLPPQALSQQHGGAPHNMNIMSHIQQQQQQQPQMQQHQQHPQSSQQQATHQQPQ

>Eurytemora affinis (crustacean; Arthropoda; Calanoida)

MLLNMEALVELPEIKSEIKAEDFAAQQQQQQHSVSVNSQQNCQWNQFGYGGEELWSSVYPGTPNSQHLVPGTPNSHLGSVTPNSHLGSVTPTSHLGSVTPTSHHAPPCTTPQTHHTLTNVITNNNAYFSQDPGLLAGYSETSAPVDLSAPRPNIHTDRYSAVDDWSGDKYNYGRYGMLPGSLTPPDKLNGDHCSPGLGPLPGSHWGGNLGLSSMAGPMQTPPSSPPMAIDRFGSSLYRSKEQRLTRDAMKKYLRERGDMVVVMLHAKVAQKSYGNEKRFFCPPPCIYLYGDGWRRKKEEMQRSGESEQGSQLCAFIGIGNSDQDMQQLDLNGKHYCAAKTLFISDSDKRKHFLLSLKMFYGNGTDIGVFHSKRIKVISKPSKKKQSLKNADLCIASGTKVALFNRLRSQTVSTRYLHVENGNFHASSVQWGAFTIHLLDDNESESEEFTVRDGYIHYGSSVKLVCSITGMALPRLIIRKVDKQMAHLDADDPVSQLHKCAFYMKDTDRMYLCLSQERIIQFQATPCPKEPNKEMLNDGANWTIISTDKVEYQFYEGMGPVRNPVTPVPVVHSLHLNGGGDVAMLELTGECFTPNLRVWFGDVESETMYRCQESMLCVVPDISQFRGGWQWVRQPTQVPVSLVRNDGIIYATGLTFTYTPEPGPRSHCTSADDIMRQGQNVDYNPASQVLM

>Triops cancriformis (shrimp; Arthropoda; Notostraca)

MPDAVPMDHLPPSARSPHDSWGQYGYGLSGYEAQQSRPDSDAGNPHSHSSTSNNEVELWNSSHSSQHHNPSSDPLSPYQGGTPLSPAGTPQQHHAYYSQNPGVVPSMTELGGSTNASLHSLTPHYLPALNNHPGGNPPSFHPEARPVDLSSSRLLAAGHAPHVLQQLGNPGTPLYLDTYNRDNRRDAYKQNGLSLSLGIGESEWQSPDPKTQNYLRYHSSSLTLTPPDKVNVDGSNSQSGGGGSQQQQLHSASSSYSIAVANMVPTLQTPPSPLSTPSPPVPLERYGGPPLFRPKEQRLTREAMMRYLQERGDMVLVILHAKVAQKSYGNEKRFFCPPPCIYLYGDGWRRRREQLQREATAAGASAAEAEASSQLCAFIGIGNSDQDMQQLDLNGKHYCAAKTLFISDSDKRKHFMLSVKMFYGNGQDIGVFHSKRIKVISKPSKKKQSLKNADLCIASGTRVALFNRLRSQTVSTRYLHVENNNFHASSTQWGAFTIHLLDDSESESEEFTVRDGYIHYGSTVKLVCSVTGMALPRLIIRKVDKQMAVLDADDPVSQLHKCAFYMKDTERMYLCLSQERIIQFQATPCPKEPNKEMINDGAAWTIISTDKAEYQFFEGMGPVRSPVTPVPIVHSLHLNGGGEVAMLELTGENLTPNLKVWFGDVEAETMFRCEESMLCVVPDISAFRSGWQWVRHPTQVPVSLVRSDGVIYATRLTFTYTPEPGPRPHCPTTDEVLRGSVSTNGSSHMVGGNMHYEHLPQGPTM

>Argulus siamensis (louse; Arthropoda; Arguloida)

MFDSNQGQYSYSLNSGYDNRGTPPEGVWHNVNGGESSYYAANSSESVPTTTGYYIKEENDDVSSNMTELGSTPAPHPPPQVQPSMHHHMNGHAPGGRYPENNPVDLSNSRTGLPELVYRDRRDIGQASHYKPNGLGLNVGIPTDSDWQNPEGKNHYGYPRYPAASPMIPGSLTPPDKMNGEHHPGHPNHHHPAPHHMMGIPTSSPLSISTMVSSMQTPPSPLPTPSPPVPLERFGLNCKDQRLTREAMKKYLRERGDMTIMILHAKVAQKSYGNEKRFFCPPPCIYLFGEGWKRKREQMLREGENEQGAQLCAFIGIGSSDQDMQQLDLSGKNYCAAKTLYISDSDKRKHFFLSVKMFYGNGHDIGVFESKRIKVISKPSKKKQSLKNADLCIASGTKVALFNRLRSQTVSTRYLHVENGNFHASSTQWGAFTIHLLDENESESEEFTVRDGYIHYGSTVKLVCSVTGMALPRLIIRKVDKQMAQLDADDPVSQLHKCSFYMKDSERMYLCLSQERIIQFQATPCPKEPNKEMLNDGAAWTIISTDKAEYQFFEGMGPVRAPVTPVPTVQSLHLNGGGDVAMLELKGENFTPNLRVWFGDVEADTMFRCQEGLLCVVPDISAFRGGWQWVRQPTQVPVSLVRHDGIIYATGLTFTYTPEPGPRNHCPQSLDVLRPGSISRNNIGQPDPAMGYDQIPPSHHPL

>Strigamia maritima (centipede; Arthropoda; Geophilomorpha)

MNEVFLPQGQYDYPPPLASTYSREADLWNVNLATYSSAPTTCTGATPAPSVTGFYAQATGSNSVSPSSVSLTTLTPHFADNHPVDLSNSHRGEGGHLDLVRFQSDRVDAYKHANGLSVHIPDHHDATSHMIAGSLTPPDKVNGEHGHQLVTMSNASQMSLGSIASSLQTPPSPIPTPSPPVPLDGVRHSHKDQRLTREAMKRYLRERGDQVLVILHAKVAQKSYGNEKRFFCPPPCIYLFGDGWRRKREQMLHEGETEQGAQLCAFIGIGNSDQDMQQLDLNGKNYCAAKTLYISDSDKRKHFMLTVKMFYGNGEDIGVFHSKRIKVISKPSKKKQSLKNADLCIASGTKVALFNRLRSQTVSTRYLHVENGTFHASSTQWGAFTIHLLDDNESESEEFTVRDGYIHYGSTVKLVCSVTGMALPRLIIRKVDKQTALLDADDPVSQLHKCAFYMKDTERMYLCLSQERIIQFQATPCPKEPNKEMINDGASWTIISTDKAEYTFCEGMGPVRTLVTPVPVVHSLHLNGGGDVAMLELTGENFTPTLRVWFGDVEAETMYRCQESMLCVVPDISAFRGGWQFVRQPTQVPVSLVRSDGIIYATGLTFTYTPEPGPRPHCPATDHILRGGSQSGLDRLPTSPDPALTYNPPHHALTPI

>Metaseiulus occidentalis (mite; Arthropoda; Mesostigmata)

MNDIYNSHHHLGEEAPYPTPYPTPYPTPYPQHAPVSIQNNNYDSSAFSASTSMAGSSLNPSPLYSDPRDHHLAMLDSYGADRKPLDMSAAHRSHPYFSAAHSPMHYRGHHPPPAHPEEPPMMIASLAGSNKDLNHKQHGLLGPPKLTGGHLPPSPLPSPPVNALARRKDQPLTRDVMARYLIERNSNDMVLVILHAKVAQKSYGNEKRFFCPPPCIYLFGDGWQRKQDQIMREYGDNTEAGHQASQLCAFIGIGNSDQDMQPLDFNGKKYCAAKTLFISDSDKRKHFMLSVKMFYGNGEDLGVFHSKRIKVISKPSKKKQSLKNADLCIASGTKVALFNRLRSQTVSTRYLHVENDNFHASSTQWGAFEIHLLDDEESESEEFNVRDGYIHYGSTVKLVCSVTGMAMPRLVIRKVDKQTALLDADDPVSQLHKCAFYMKDSERMYLCLSQEKIIPFQATQCPKEPNKEMINDGAAWTIISTDKAEYTFFEGMGPVRAPVTPVPVVNSLHLNGGGEIAMLELVGENLTPDLKVWFGNVEAPETMFRCEESMLCVVPDISAFKEGWQWVRQPTQVPVSLVRSDGVIYATSLTFVYTPEVGPRQPYPCVGDILRPQSQLQSVSPGCGQQQQASQLQDDVNSQQSQSSTNHYMHLQ

>Ixodes scapularis (tick; Arthropoda; Ixodida)

MSDVYLPDDQYAGYHPVQNNYSSPHDDGTFAVSGSAYSGDYGRDLLLDLGQQAAGAPVDMSSHPARAHPYFNSGGVPFKNGLADGEPGALLGSAAKGAGEGPPQGSPHLGVQAPRLPPSPLPSPPSEERYRRGEPRLTRDAMDRYLRDRGDMVLVILHAKVAQKSYGNEKRFFCPPPCVYLLGDGWQRKRDQLLRDGEADQAAQLCAFIGIGNSDQDMQQLDFAGKSYCAAKTLFISDSDKRKHFMLSVKLFYGNGEDVGVFQSKRIKVISKPSKKKQSLKNADLCIASGTRVALFNRLRSQTVSTRYLHVDGGNFHASSSQWGAFTIHLLDDNESEAEEFTVRDGYIHYGSTVKLVCSVTGMALPRLVIRKVDKQNAFLDADDPVSQLHKCAFYMKDTERMYLCLSQEKIIQFQATPCPKEPNREMINDGASWTIISTDKAEYTFHEGAGPVRVPVTPVPVVNSLHLNGGGDIAMLELTGENFAPNLRVWFGNVEAETMYRCAECLLCVVPDISAFREGWQWVRQPTQVPVSLVRSDGVIYATGLTFTYTPEPGPRQPGPYPALVHDILRPANARSHPPAPEDHGAPTANFGHHMHFGSHHQNMS

>Parasteatoda tepidariorum 1 (spider; Arthropoda; Araneae)

MSLAPYTTAYSSASSTPSQNASSYSITSQANLQNPNPYTDLNVNNRAQHQQLGHVQLNEVNRNGMQNSHLCSPSRDRPSENVIDSHPVDLSSPKPSNRYGPMGYMNVARDNSTGAFPNFNNRSGFENGMDSMRPEYNQKLNGEQNQQQPHHPLVPHGALSLPLLSQGRGHAPPSPMPTPSPPVDDRHRSKRKDQRLTRECMKKYLRERGDMVLVILHAKVAQKSYGNEKRFFCPPPCIYLLGDGWRKKQEQMVRDGESEQGSQLCAFIGIGNSDQEMQQLDFNGKASTQYSYCAAKTLYISDSDKRKHFMLSVKMFYGNGEDIGVFHSKRIKVISKPSKKKQSIKNADLCIASGSRVALFNRLRSQTVSTRYLHVENGNFHASSTQWGAFTIHLLDDTESESEEFTVRDGYIHYGSTVKLVCSVTGMALPRLIIRKVDKQTALLDADDPVSQLHKCAFYMKDTERMYLCLSQERIIQFQATPCPKEPNKEMINDGASWTIISTDKAEYTFYEGMSPVRSTVTPVPVVHSLQVNGGGDVAMLELTGENFTPTLKVWFGEVEAETMYRCQENMLCVVPDISAFREGWQWVRQPTQVPVSLVRNDGIIYSTGLTFTYTPEPGPRTHCPAVEEILRPPEMRRDDPRHPSHMYSNADHPMQ

>Limulus polyphemus 1 (horseshoe crab; Arthropoda; Xiphosura)

MIEIKRERFESCQEVDVGVMGNNHAYMTEDQYGYTLGSGYEPPPNSTLPLTEATLLSHYSTGNTGPSSPGVNNASVYSTSRGGGICQQGSIMMDISDSSVMNGDVVGHHASDGTFQHLQSRVRANDHSGRGPYDGHPVDLSNQRPDSHLSHMNLGSYVGTHYRSMSQNRSHFENEHSVSENESSMQMLSSSLSVSEKPTIDHQQVNHLPPFSGQNPLGGSTLGSRTHPSPSPLPTPPPIDDDGHRSRRRDQRLTREAMKNYLKERGDMVLVILHAKVAQKSYGNEKRFFCPPPCIYLLGDGWRKKQQQMVRDGENDQSAQLCAFIGIGNSDQEMQQLDFNGKVWKNYCAAKTLYISDSDKRKHFMLSVKMFYGNGEDIGVFHSKRIKVISKPSKKKQSLKNADLCIASGTKIALFNRLRSQTVSTRYLHVENGNFHASSTQWGAFTIHLLDDNESESEEFTVRDGYIHYGSTVKLVCSVTGMALPRLVVRKVDKQTALLDASDPVSQLHKCSFFMKDTERMYLCLSQERIIQFQATPCPKEPFKEMINDGASWTIISTDKAEYTFYEGMGPVHSPVTPVPVVFSLHLNGGGDVAMLELTGENFTPVLKVWFGDVEAETMYRCQESMLCVVPDISAFREGWQWVHHPTQVPVSLVRNDGIIYATGLTFTYTPEPGPRPHCRTVEEILRPAGSHTHPPDGEEPGPLISTYTHVSSGL

>Centruroides sculpturatus (scorpion; Arthropoda; Scorpiones)

MNDVGIFQTEEPYGYPVTHNYQEENRNFDVSSSVQIPQYSVVISSPITTPSSNASVYSLPIQDSNSHRQNMMMELGQSQHSQQSVNHMQINGDVLGHHNSNSLLHSQTQHNASSEHRGSGYESSPVDLSSHRSVGRFGPVDMDAHLGAASIAHYRNLSQEDNSFENGLRNAVPVSMSDTESSMQLISGSMTSHDKVNGDQHSLGSLNSSNALSVPMLAHGRGHAPPSPLPTPSPPVDDRHRNKRKDQRLTREAMKKYLRERGDMILVILHAKVAQKSYGNEKRFFCPPPCIYLLGDGWRKKQEQMIRDGEIEQGAQLCAFIGIGNSDQEMQQLDFNGKNYCAAKTLYISDSDKRKHFMLSVKMFYGNGEDIGVFHSKRIKVISKPSKKKQSLKNADLCIASGTKVALFNRLRSQTVSTRYLHVENGNFHASSTQWGAFTIHLLDDNESESEEFTVRDGYIHYGSTVKLVCSVTGMALPRLIIRKVDKQTALLDADDPVSQLHKCAFYMKDTERMYLCLSQERIIQFHATPCPKELNKEMINDGASWTIISTDKAEYTFYEGMGAVKTPVTPVPVVHSLHLNGGGDVAMLELAGENFTPNLKVWFGDVEAETMYRCQEGMLCVVPDISAFREGWQWVRQPTQVPVSLVRSDGIIYATGLTFTYTPEPGPRPHCPGVEEILRPAGLHQVDENQSRVTPNTVPVAPPHGYPEAHM

>Euperipatoides kanangrensis (velvet worm; Onychophora)

MNEVYISQGQYGYTVGANFQRDNEILYGLNGIPTSLQYPLQASSHNDQQTQFFLQDGSSEQQQQQHNLSDMNRTQQHAHINGHIQPNSYDNPVDLSSHRSSGGGNGRGHVEINGHLNKHPHHHGTVYRDPPPAHNGTYKQHNGLNIPDHIESSQQILPGSLGPSDKVNGDLVSLATTSPLSISTMATTIQTPPSPLPTPSPPVEDRRHGHKDQRLTREAMKRYLRDRGDQILVILHAKVAQKSYGNEKRFFCPPPCIYLFGDGWRRKKEQMERDGETEQGSQLCAFIGIGNSDQDMQQLDLNGKNYCAAKTLYISDSDKRKHFMLSVKMFYGNGEDIGVFHSKRIKVISKPSKKKQSLKNADLCIASGTKVALFNRLRSQTVSTRYLHVENGNFHASSTQWGAFTIHLLDDNESESEEFTVRDGYIHYGSTVKLVCSVTGMALPRLVIRKVDKQTALLDADDPVSQLHKCAFYMKDTERMYLCLSQERIIQFQATPCPKEPNKEMINDGASWTIISTDKAEYTFFEGMGPVKAPVTPVPVVHSLHLNGGGDVAMLELSGENFTPSLRVWFGDVEAETMYRCEESMLCVVPDISAFRGGWQWVRQPTQVPVSLVRNDGIIYATGLTFTYTPEPGPRPHCTAADQILRTTSTVSTVSYNSM

>Octopus bimaculoides (octopus; Mollusca; Octopoda)

MNEKDHMNTSQNSFLYAFGNSTETGAQLMYTSSNHGLPGSVCTSEAGYIVYTTAAPGMESVTDSSGLLPQPHLQEVITEHLDNHLVHPRTTLINGRMAQNGFDNPMDLSNGKVVQVIDMKKEDSRQYDHNQAATNYHHNGVTVGIPESQHGASEHLMPAGSLTPPDKISGDSISMANIRPLGISTLTNTIKTPPSPMPTPSPPINRITGDIDHSDSRITHPFSRSHWTTDVLENRHVGQYQDQRLTREAMRKYLRDRGDQVLVILHAKVAQKSYGNEKRFFCPPPCIYLFNNGWKRKKEQLERDGATEQESLVCAFMGIGNSDQDMVQLNLEGKNYCAAKTLYISDSDKRKHFMLSVKMFYGNSQDIGMFNSKRIKVISKPSKKKQSLKNADLCIASGTKVALFNRLRSQTVSTRYLHVEDGNFHASSTQWGAFTIHLLDDNESESEEFTVRDGYIHYGSTIKLVCSVTGMALPRLIIRKVDKQTALLDADDPVSQLHKCAFYMKDTERMYLCLSQERIIQFQATPCPKEPNKEMINDGASWTIISTDKAEYTFFEGMGPVKATVTPVPVVSSLHLNGGGDVAMLELTGENFTPILKVWFGDVEAETMYRCEESMLCVVPDISAFRAGWRWVRQPLQVPVTLVRNDGIIYATGLTFTYTPEPGPRQHCNAVDRVLRANSDPPPPSSVNYSAPPM

>Crassostrea gigas (oyster; Mollusca; Ostreoida)

MNEQHLYVSQNGYTYSVGPTQQSSHLMSCSQPQQNGSHTGHGFMHHNAGYTLHPPDRQYGGRADPGPSSRADLTGHMAHGGYENPMDLSSNKPGNPGRLVKEEGHHHGYLAAMGTTPVSVGIPDHVHNPSHIVAGSLTPPEKINGDPGAMATSSPLSITTMTQAIPAPPSPISTPSPLYASNSYVDRGYQDQRLTKEAMRNYLKDRGDQVLVILHAKVAQKSYGNEKRFFCPPPCIYLFGSGWKRKKEAIEAEGGTEQDSTTCAFMGIGNSDQEMVQLNLEGKNYCAAKTLYISDSDKRKHFMLTVKMFFGNGQDIGVFNSKRIKVISKPSKKKQSLKNADLCIASGTKVALFNRLRSQTVSTRYLHVENGNFHASSTQWGAFTIHLLDDNESESEEFTVRDGYIHYGSTIKLVCSVTGMALPRLIIRKVDKQTAILDADDPVSQLHKCAFYMKDSERMYLCLSQERIIQFQATPCPKEPNKEMINDGASWTIISTDKAEYTFFEGMGPVKSPVSPVPVVNSLHLNGGGDVAMLELSGEFLAPNLKVWFGEVEAETMFRCEESMFCVVPDISAFRAGWRWVRQPLQVPVLLVRSDGIIYSTGLTFTYTPEPGPRAHSREVDRIIQPGVSSPDSTSTANFSNPL

>Lottia gigantea (limpet; Mollusca; Docoglossa)

MNEHHLFGSQDGYGVNLDGANLSHHGVHYAIHSNGNFVSSQQGRGIKRTNHELPPHLNTVHYLINGHMAQAGVENPVDLSNGRISMTGLSSDSDNQYERHSDDRQSSQQQQSQYRHSGVRVGIPDQSHDATSHLFTGSLTPPEKPNGDLVPMSTTSPLSITTIGPPMQAPPSPLPTPSPPMRHPPDGSGLRVPQCDAPHLSMRKDQRLTKEAMRKYLRDRSDQILVILHAKVAQKSYGNEKRFFCPPPCIYLFGKGWKRKHDQMEEEGSTKDEAQVCAFMGIGNSDQEMVQLHLEDKDYCAAKTLYISDSDKRKHFMLSVKMFYGNGQDIGLFLGKRIKVISKPSKKKQSLKNAELCIASGTKVALFNRLRSQTVSTRYLHVENYNGKCNFHASSTQWGAFTIHLLDDNEGESEEFTVRDGYIHYGSTVKLVCSVTGMALPRLVIRKVDKQTALLDADDPVSQLHKCAFFLKDTERMYLCLSQERIIQFQATPCPKEPNKEMINDGASWTIISTDKAEYTFYEGMGPVKNSLTPVPVVNSLHLNGGGDVAMLELNGENFNPSLKVWFGDVEAETMFRAEDSMLCVVPDIAAFRPGWKWVRQPLQAPVSLVRLDGVIYATGLTFTYTPEPGPRQHCKDMDRIVGRTSTASPDSTSHTTL

>Notospermus geniculatus (ribbon worm; Nemertea; Heteronemertea)

MRSVDKIMNEQEIYVSQGQYGYTIGVNQQRDNRHLYPNNVMPCSLQAADTNGLLPENTKTMQNGNMSDQLVSHNRMHPNGRMAVQYDNPIDLSNRLEGATQHAQGRDLNGYGKIPRYEHHVDTTSYRHAHSQNQSGQNKVTSSMNMVGIADQSHESGNAHNHMIPGSLTPPDKVNGDMVPLPTTSPLSISTMATSIQTPPSPMPTPSPPLSHHRSEHDTRHHNGQFQDQRLTRDAMRKYLRDRGDQVLVILHAKVAQKSYGNEKRFFCPPPCIYLFGNGWKRKKDQMERDGAAEQESQVCAFMGIGNSDQEMVQLNLDGKNYCAAKTLYISDSDKRKHFMLSVKMFYGNGQDIGVFNSKRIKVISKPSKKKQSLKNADLCIASGTKVALFNRLRSQTVSTRYLHVEGGNFHASSTQWGAFTIHLLDDNESESEEFTVRDGYIHYGSTVKLVCSVTGMALPRLIIRKVDKQTALLDADDPVSQLHKCAFFMKDTERMYLCLSQERIIQFQATPCPKEPNKEMINDGASWTIISTDKAEYTFFEGMGPVKSPVTPVPVVNSLHLNGGGDVAMLELSGENFTPTLRVWFGDVEAETMYRCEESMLCVVPDISAFRAGWRWVRQPLQVPVSLVRNDGVIYATGLTFTYTPEPGPRPHCRDADQIMRPGGTVQALQSPDSSSTHFVPVPNQM

>Malacobdella grossa (ribbon worm; Nemertea; Bdellonemertea)

MNCELYVPQGSYGGYTIGANLQRDIPLYQHYNNYLNQQQQPQQQPARPENNNYRQQIYGNGYNANQQLQQPTGQQSSTDMVDSPFCLSDNANKQNGSHFEAKRNANGNPYNAKMASERLSIPSALTRALHYDNPLDLTNRLDEVREDARHDCAKIARLAYDDDISCARKNSPSALTTLHAGLPDSSHDGSGIAGSMTPPDGGKGNDLDLQSSMATTSPLSIQTIANSIQTPPSPVQTPSPPPPLLRSCRNENDGHNRNGLQMAALPRGSHYGHYSDQKLTREAMRKYLKERNDQILVILHAKVAQKSYGNEKRFFCPPPCIYLFGRGWKRKRELMTSDGATDQESLVCAFMGIGNSDQEMVQLNLEGKNYCAAKTLYISDSDKRKHFMLTVKMFYGNGQDIGVFHGKRIKVISKPSKKKQSLKNADLCIASGTKVALFNRLRSQTVSTRYLHVDGGNFHASSTQWGAFTIHLLDDNESESEEFTVRDGYIHYGATVKLVCSVTGMALPRLIVRKVDKQTALLDADDPVSQLHKCSFHMKDSERMYLCLSQERIIQFQATPCPKEPNKEMINDGASWTIISTDKAEYNFYEGMGPVKAPVTPVPVVNSLHLNGGGDVAMLELTGENFSPNLRVWFKDVEAETMYRCVDTMLCVVPDISAFRAGWRWVMEPLQVPVSLVRSDGVIYSTGLTFTYTPEPGPRPTWRDENSLHANTTLCNGGLTNQHF

>Lingula anatina (brachiopod; Brachiopoda; Lingulida)

MNEAEIYISQGRYGYTIGANLNRDDHIHSYLDSQQQQHVACANIIAEELIYQATDSKFTEDGYVKMNRETASQNGRRGGYENPMDLSRRTEMNGHVGHSYEQRDQSYINHNSVSSSVSVAIPDQSHEASSSAHMIPGNLTPPDKVNGEMVPMATASPLSISTMATSIHTPPSPLPTPSPPVRHPGDGERGDMRHINGHFSGQRLNREAMKKYLRDRGDQTLVILHAKVAQKSYGNEKRFFCPPPCIYLFGSGWKRKKEQIEKEGGSEQDSAVCAFMGIGNSDQEMVQLNLDGKHYCAAKTLFISDSDKRKHFMLTVKMFYGNGQDVGVFHSKRIKVISKPSKKKQSLKNADLCIASGTKVALFNRLRSQTVSTRYLHVENGNFHASSTQWGAFTIHLLDDNESESEEFTVRDGYIHYGATVKLVCSVTGMALPRLVIRKVDKQTALLDADDPVSQLHKCAFYMKDTERMYLCLSQERIIQFQATPCPKEPNKEMINDGASWTIISTDKAEYTFYEGMGPVKAPVTPVPLVHNLYLNGGGDVAMLELSGENFTPHLKVWFGDVEAETMYRCEEGMLCVVPDISAFRSGWKYVRQPLQVPVSLVREDGVIYATGLTFTYTPEPGPRPHCAAIDETMRPGASVIQTTDSVHETQHQQYSNSSSTAVS

>Phoronis australis (phoronid worm; Phoronida)

MTTMATQGQYRYSIQRDTYASHPAFDQDAFCPDPADFECFRTGTLDVSDVRQTHLSVVDSMALSSRQHDNRPMDLSSRGQHHHHHHHHQGVLSSVHHRALKDHYTAEQAAMFYGNVTSSVTVGIPDQSHESASHLIAGSLTPPDKVNGDVVSMATASPLSIGTMAPSIQTPPSPLPTPSPPLSRTEMDSGYNGHFQEKRLTKDAMKKYLRERCDQTLVVLHAKVAQKSYGNEKRFFCPPPCIYLFGSGWKRKKEEMEREGRSEQESQVCAFMGIGNSDLEMVQLNLDGKNYCAAKTLYISDQDKRKHFMLSVKMFFGNGHDLGVFNGKRIKVISKPSKKKQSLKNADLCIASGTKVALFNRLRSQTVSTRYLHVEDGNFHASSTQWGAFTIHLLDDNESESEEFTVRDGYIHYGSTIKLVCSVTGMALPRLIIRKVDKQTALLDADDPVSQLHKCAFYMKDTERMYLCLSQERIIQFQATPCPKEANKEMINDGASWTIISTDKAEYKFYEGMGPVQNPVTPVPIVHQLHLNGGGDVAMLELSGESFTAKLKCWFGDVEAETMYRCEESMLCVVPDISQFRAGWRYVRQPLQVPVSLVRNDGVIYATGMTFTYTPEPGSRPRCPASEAMLHGSVTMASPDSTYNNCGS

>Xenoturbella bocki (paradox worm; Xenacoelomorpha)

MSTHLVPTTTFDYGFSTNHSQLPSSHYAAVYHGLPSTERIPLDRHSTDPPGDGELTPRFAEVFHTVPTTGSDVYGSQKLINPQSCGMPQLYPSNINSHLLTNDNHRRMSCDNNGSHMTVPQDGMYDDRTNGHELKTSPAVVAGAVAMETSAMTRKQHVFRAPRHLVLPVATTYYESPPTTKKSRTVEYSMRSPLEGATNTSPYGMMGHAPQGLGGHSPSLGGHSPALGGHSPVLGEPADGDLPVLKSSELSSMGAKRYSAPLNLTVHDKCKDVRVLGRLTPPDKQHVNNDVGALLSPSALSPHSSDASDTESTQPPSQTIPLTPPIDDTQYSCSKHLTREAMQRYLNDRPDCTLLILHAKVAQKSYGSEKRFFCPPPCVYLLSSGWKIKRDHTAESQICAYMGIGNSDQEPQLLNLDNKTYCAAKTLYISDSDKRKHFMLSVKMAMSSSGLGADIGTFMSKRIKVISKPSKKKQSLKNTDLCIASGTKVALFNRLRSQTVSTRYLHVEDGNFHASSTQWGAFYIHLLDDAESESEEFTVREGYIHYGSTVKLVCSVTGMALPRLIIRKVDKQTALLDADDPVSQLHKCAFFMKDTERMYLCLSQERIIQFQATPCPKEPKKEMINDGASWTIISTDSAEYSFYEGMGPVTRSVTPVPIVHSLQLNGGGDVAMLELSGESFSPAVKVWFGDVEAETMFRCTESLLCVVPDISAFRGEWRWVRQPTQVPVTLVREDGIIYSTGLTFTYTPEPGPRHGSITDTLRLHTTANTNTETTTTDHYINPCNSYPGL
