## Supplementary file 5 for "Hairless as a novel component of the Notch signaling pathway"

**Supplementary file 5.** **Full-length S-CAP sequences**

>Strigamia maritima (centipede; Geophilomorpha)

MPNPNCETCESSQNNKRKCENNANVQSGNKKRSTDPNECQQPAQSAHDFLKKLVHEQKLHAPTLFNNTNELKPENNRTGSSENDTNGKLSFYLDGRFILELEHKPGGKRRNGGWIQTKGITWMPRPEEEKLINPIKQELKESETSRMKLKNLKRFSSQRETTNACSVVLFNFRINRRRDFKHFPRLNAYYLRKAARHPFSIDLFFFSKSEKCKTAPPSTKLVKKRTKRLNDLFRPPEFVNSTADNVKRWFDEVGSVKVKCEPDNASLPLPEWYLDPSKSQIPVNEMIGNPRPYLPAAPTHYDDKESNVNSMSEPLDLRTNVVRETKRDEWMKREAKEHPGNAFYVGLIPFPLPGHGEYSAPSAVQPWPCVSCPVVAYAPTVVNSYYYCCCFLPGCRSNCLCKTEGKQLTSRSSEIPKECDCATDLTNACNKKFPIPVAVKSDHQTPGKKTDVGDGCSHSMLKDLLLMPSSEKDCHCPQTSNP

>Nephila clavipes (spider; Araneae)

MDKCSTDAEFDSYWDYEAKNKNRNRIRVGRQYQATIPPLLKPGQKDGRKSEELETLKWKPDQLSDQALEEYLSMAKGIGLFSKSLDSPKPSEKSDNSLQSAIKGLSEFVTSHHPCHHDDGCKVARPSSSGESSSKSTATSDWSPSEAQLFAQALEACGKNFGAIKKEYLPWKPIKSIIEYYYEGKNEKLEASSEAETSGKISPKKELGSPASYKEEKDFIKEEVLPDEAVPEGSIPEDPPKDAINFLENTRVTDPCRATPVELKMSVNENDIVTASSTVASLKFFLGGRLVLKLNAQQDSGSGTKCQWVQSNDLPKHSNNIKKDRQKKKCIQDESFVINPDVHKLKKKTEELSSASNELTDCDPSAVKKAKLKSEYDFPLCSSSWTPPVADGQESIEVDDASSKTSEGYISPVLSNSRFSKADAKLDSVSNFNSVGDPKCDSDMCWVKSEMKGCNISSDHCNDSASKNICMLGGKNSPNHMSDVSPYKQEVLNHKSALSHKTCVKSLCYQPHTSHSNRSLSKQWSKRSNMTYPMRTSPCSLSPISVVDLKSPSKETPLDLSSKLDNVLNDIDHHNSQSAHKTKVQVDNAEKIMPSESESRPASSEFSPQSSNHGSLAQSIDSPYKDTINSHKCSEDTASDHQQCEKCLRQEDSDFSSGKDSENSCQLPYTYCNLPYQCALPKCTKDCLLEEHLCNKSDAEKSQAGSPGYIINDVDENFCKKNPAWCSRSGASYYPYYSQCYPYYLSYGVAPSHVSEPTIAEVPLDLSSTNDKVKENSSETEKLPEEVNVMNSDDTDAVERGMLYQLLKNKTSK

>Parasteatoda tepidariorum (spider; Araneae)

MDKCSTDAEFDSYWDNEAKNQNRNRIRVGRQYQATIPPLLKPGEKDGRKLEDLETLKWKPDQLSDQKLEEYMSMAKGISLFSKSVSSSRSPEKSDNSLQSAIKGLTEFVASHHPCHHDDGCRQVLKPSSSSTESVSKVNTNTWSQSEAQLFAQALEACGKNFSAIKKEFLPWKPVRSIIEYYYQIKNEKAEETEPTDKVIPKAEEEPIPSCSHEEKPCVKEEPVKVEEPQVKTVNTEDPPKDSINFLDHSRVTDPCSAASETSLPQDVPATSTVGSLKFFLGGRLVLKLNAQQDGGSGNKCQWVQSNDLPKHSNHNKKDKHKKKFAPYSYSSSGTQKPLKKGDDTSAVPDCDPSGIKKPRLKEYETSENSALGLLLCSSSWTPPVADGQESIDVDDTSSKTSEGYISPILSNNSRTSKIDTIKHDFASNPNTSNELGEQKCDSDPTWVKTDNCIISDSASKNASVLSGKNSPNQCYSSPMKKEVLNHKSALSHGRSVKSPCYQSRSSHSSSPLLKQRNIKWSNIPYPLKSSPSSLSPVYIDLRSSTKESAIDLSSKSDNGLNNADHYVKQTHSNERSECNECILCESDSRPDSEFSPQSSNFTPSAESPRKEPSCTACNCCEGVSTDQLPFDNSGRKEDETMTNKESDDNSQLSYTYCNLPYHCALPKCTKDYTLEEHLCNKSSINKNQVDCPEYIINDIDENMCEKNPNWCSKASYIPYYSHCYPYYVSYGVTSRNSAESTDTEVSEAPLDLSSTNETKPEKLPNVEKISENSSEEVSQNAEGGRGMLYQLLKNKSK

>Stegodyphus mimosarum (spider; Araneae)

MDKCSSDAEFDSYWDYEAKNKNRNRIRVGRQYQATVPPLLKPGEKDGRKCEELETLKWKPGQLSEQEVEQYLSMAKGIGLFAKAVDSPKIPEKSDSSLQSAIKGLSEFVASHHPCHHDDGCQVPKPSSSGETDSKVNATNDWSPSEAQLFAQALEACGKNFCAIKKDFLPWKPVRSIIEYYYHGKGEKTENLADDSDTKLPEKKEEDEPVASCSHTEKSSESEMLPSDEPQSEDLNVEEPTEPTKESVLFLENSRVIDHCQAVPELKMSVTEDGMPASSTVGSLKFFLGGRLVLKLNARQDSGSGSKCQWVQSNDLPKHSNTTKKVKHKKKSMQDGSFPASDIFKQQKKSEELCISNRVSDYDTSVVKKPKLKLEHISHLDKQELLLCSSSWMPPVADGHESVEIDDGSSKTSEGNGNVSPVLSNPRSIKADVKTDFASNFVNELGEQNCDSDPCWVKLESQCGNVATESSFVDNSASKCVSYVGENNSPNQFIDDSSFKQEVLNMSALSHNKAHTKNQCYQPLTSHSNSPLLKQWSKSPNPVKTSPSSLSPVSLIDTKSPVKDIPLDLSSKLENSINSTDHHDMMDIHKPTVKVECSNQSFPSETETSPVNCEFSPQSSKLGSHVQSNESWRKETKNNVYQNSEDKADPLQYVKEENNEYSPKKRCGDPSQLPYYCNLPYHCASPKCTKDYSLEEHLCNKSNVEKNQENDPSYILQNVDENVCKKSPWCSGTVASYLQLYPPCYRSYYYSYGMAPSPASEPTIAEVPLDLSSAGESIAEKIAMPENLPEDITDVSQTNTEAVERGMLYQLLKNKTSK

>Achipteria coleoptrata (mite; Oribatida)

MDVSNSDADFDSYWEEEARNPNRNRIRIGRQYQATVPPILKQGESDGRKLEELETLRWKPDNELTDQQLDQYLSVAKAVSLFTRAIDNNYSQSNVNSDSHIENDETNETNSNSNQNTNRDKTENKNDKNKDNTLSPGSTSTVKDNRIQSALKGLSDFVSSHHPNHRDIGCRTPLLPETTANSAESESANSLTNLLSAEWNSNEIKLFSRALEVCGKNFGAIKKDFLPWKSVTSIIERYYLGIGRDGPNSGIKHEEQNNINQSGNSQNNKQESVPNTFVSDILSSVSKFMTSYTDAKPELNECQRTQSSQMSSEDIKDCDLKNNFNSEDSKIASVSGQEVKPLKAKPILPTAIQESNNSVSTVGSLKFYLDGQLVLKLNAQQEMMGQKCQWVESSDTAKKARLLHKTKKRLLNERHDSDVTNHSNHKNSNSEELDDGSNESSDDDSMESNESALVPSPSAFIAKKAKVKVENSSHKPLSSPTSLPMKDSKLDVSNRNVKIECSNSSVKKEVHLSPNSNHCEDEHKKRFSSEHTNERTNSQNCLENKWYHTIVEKSSLQPPKAHSMTSSGPLYRHSRYDSVQSSLGSSSSTSSPSSSSSSGTGVPLMSSNAIPVDLTRKSTNYSPFEAKSSSLSSSSTPDFGNYPERDKNSKVLSSPSPPRLQPPFVFPYPAYLPLSPTQSQNKIKREKEVKSSNNLCQKAVEGRPLVKSSPPTSPSLMAPSSTPPLPSALPSPSQLAWIQNSFAPYLPIYEQYYRYYYGYGMPSPNHSPGERMSSSQTPKENEANSKERAKCLTDAVLVDGDDANCGS

>Sarcoptes scabiei (mite; Astigmata)

MLDIDFDSYWDEEARNPNRNRIRIGRQYQAQCPALLKSGQNDGRKLEDLETLTWNPNNPLNIHQLNQYFSIAKSIGLFIQAIDTCQDGIDEIECLEEEKCSTRPFQILDSGTKCILEAKPDRTVQSSNDNNNVINTSVSIPEKHQSDKVTTRQRIQHHHKSNHKEDSVEVTSQSSSNSSSSSSTTSNSLSTSPLKQTSNTRIQSIAKGLPAFILSHHSKNMREEKITDDDDYESKNNKGKNDDELQDSKCKFKSTTAEQAINLGSVEKYDVKKLANLLMGKWNATEAKVFAQAFNQCGKNFMAIKKDYLPWKSIRSIIEYYYLTCDKEKEELRKQRRNRAKSNRNCFSLNENRTNTTKPPSSSDNSDEINYESAKKFSTDIDGSNPKFNISQNSNQKSNDDRTRLSPNKQSNLSEQDLFNLMLMKGGLKNGNINLDAIADFTNSRFKFNNTNNNCNINGGNSAFIQSSSESKNSFVPGQEVRPVKAKPIFSQTDPSSSQLNEADSSKTNLGSLNLYLHGELVLRLNAQQQDSGQKWVESNEIQSSFKDDADDFSCGGADVSASDDDSLTSNESSSLVASSPSSSSTIATSSAKKSRVKLEHQNSNFPSNSQNSSISSPVSSHRNKNESNFNGGALNNPFPNTSQSLINSISALAAASSFLENPLSELEIKKRLFEYCQMGMSVEELQLLLASNLFPDPPKAHSNTPCLAATVPHIGTNVTDKTTTQNVSEKDSVYRSSPVKEHQRSSKSSSKQSKIVSNDLDNADNGYTNGFGPIDLTRRKSSINHNPFNSISNEMGSKNHSPKSKQSFSASTSNNLLNRSNCSLFSLPTLSSSSPSSLSSSTLSNPNGTGRNSTIVSPAKFSSNSHSSKSMKNNYNSKKKSSSSSSSSSISIAETNPF

>Metaseiulus occidentalis (mite; Mesostigmata)

MDSNNDVDFDAYWEEESRNKYRNRIRVGRQYQASVPPLLRPGESDDRQLEDLETQYWKPSLDDEAIHEYLSMAKAVSVFSCSLESQSHDNLQSAIRGLTEFVMKHHDPCHRDAGCRMSVKSSWTNKEAELLACALERCENANRKKMAEEYDESGSDDADGVDETSDIEADCQPSSTSALELPIDVNPPSEIKPVPARVIGRKTDSPVSDSGSPTGPAQATGTAAGGATPVSAKSRADGANTSQQLQDPSAAGQGSLKFYLKGQLILKLNAHQERKTWVEDPDNPAASSGQFGGVAGALSGGQNRRKAGRKANRSSTTSGPISWLDRGSPSLESTSSLDSHNSTSTSATTPTCAAATPTPSTTAATSTAATNALPAGGNNGGGGGGSSSSSSSSCSSSSSSAASSAANPVSQQPAPLDLSSQQS

>Dinothrombium tinctorium (mite; Trombidiformes)

MDVSTSDADFDSYWEEEARNPNRNRIRIGRQYQATVPPLLRPGESDGRKLEDLETLKWKPENDLSDQQLDQYLSLAKAVALFARAITNNSSANQTDSESAHSNGNSGNNTNSNSENQADEEQNSTSQEDKKNAENTDSFVSQPSDHLQCALKGLSDFVTSHHPFSRDAGCKVPINSSDKSATSSSQSNSENFTNSTKLNWTSDESDIFAKALDACGKNFSAIKKDFLPWKSVQSIIAYYYLGLNRTKLDTNCSDKNIKAFSSNDEKDCTTPKMSKDCSALINVATTSTASSPNSPTFSLSPIMNTACKDYSSKKHVNGVSNSLIECDSNSKQNEANCVNLEIKALKAKPVLPSNDDLSCNMSNLGSLKFYMDGQLVLKLNAKQEVTGRKCQWVESQDTPKFSRPVRKGNKKLLLEKSESDSKIGHVHNDKCMHQSNEDGEEGSLDSSDEDSLESSESNAVLSPMSVTSRKAKVKVENNCFVPLPSPCGVSGSKDQSLNRSNSTNESPTSVVKKEPLCSPTSSSNQKNDIEAKRKLKVPPWPPDKKKSLFITATDNKWSIAEKNVKPPEAHAASRILNSGSYSSSAVPVDLTRKSSAYSPNGTKSHSSLSSSSSTSGDAEKISSSPAQLNYPTSLTPNSLSSSQKQSKSAKDLKPIEMCQKEKSPKSDKLSAEKTNSVSNASLAWFQSPLVSYLPIYEQYYQYYCRYGFPPLPNTAASCESKSNKTSNH

>Ixodes scapularis (tick; Ixodida)

MPNSNRLLARRPPCEVPLDADFDSYWENESRNNNRNRIRVGRQYQATVPALLRPGESDGRRLEDLETLRWRPESLSDQSIDEYLSMAKAVSLFARAMDKWQAWGEGGPEGCCLQTALRGLSDFVTSHHGCHHDAGCQVGPTLPCHWTPTEASLFARALDECGKNFGAIKKDFLPWKPVKSLIEFYYQGRMPKQESDGQEQEAGCSTSTGASCTSNCSTSKCTANCTSNCISNCIKNCIKKEVKEEPVEEDEEEEREEEKQKGEEPPPEGTAEEAPGVSGPEVKPMRAKPVKAAEEGPASVAPVGSLKFFLGGRLVLKLSAQEGGAWVEAQDTPRLGRPRQPPPDDASDEEEPSPPGSGGGGPPQKGTSRCTSWPETATPAPASSSAGGASASTPEENNTQNSTEEEEAGDESPPLPAPLREASPKCLPPPASAAPVTGAGTLPPPGCAERCRPLSGWRSPRGAVPFLGPLFAAQCCPTTFLPASTADAPLDLSSPVDAKKLPLVPK

>Centruroides sculpturatus (scorpion; Scorpiones)

MFRLMDASNSDADFDSYWEDQSRNRNRNRIRVGRQYQATVPPLLKPGETDGRCCEDLETLKWKPDNDLSDQQIDQYLSMASRAVSLFAKAIDTCQNTGNGPDKTLQTALRGLSDFVTSHHPCHHDAGCQVPSPSCCGDHNCKHSIKNGWTPTEAQLFARALEACGKNFGAIKKEFLPWKPVRSIIEFYYNGKEEKYDNQSARQSQQIANLSENLITGKENISEAHNLKNEDEEDDDDDDEVDKLEDCDCKLEKTDSKSDHDTETQNEKTSKLNQPNNESTSFNLGSLSFFLGGQLVLKLNAQQENGPDNQNCQWVLSQDTPKLPPNRKGRKSKKSCKQGSITSIGKEEIQQTKNEKKQEFCMEEREEGSGVKKAKLKTDCIHSSSMEEASDFCNMPWLPPTETKQERTEKCDDTLYKTETWCHSFSSLCLSQCKNENCTHKDKNCLKNTGLLNSEDNITSTIKTEPGLSPSYIMNSNVSLKEEDSKESVVSMKSYLKALSVPSSSLSSQNWVEHSNIWRNSTKFNEILTKKETWSHSQTLQDSDGGPLDLSIKNSEKNLSKSCEALTSEKSLTVHDSVASDTQMCPFIKSSTCQKRTSVSTSIKDSLSLNDMSQTNMPLDGWSVQVSHGLPYVKGETVKKDCHSSQLLSKGYPLSCEYSRYGMTVTYAPTVFPPVYNCKKDPTNKDKPVNLSKKNENFTCEENNSTRELEVDERCSESGVDLVPWVVAPILSYGQMHSPFCSCCVTYTDSMGMNRLSDQPQGDSETPLDLSSSSSSSSSSSSSSDTTDITDFHKPKNENVAD

>Limulus polyphemus (horseshoe crab; Xiphosura)

MDASSDADFDSYWEDESKSKNRNRIRVGRQYQATVPQLLKPGESDGRQLEDLETLKWKPNNDLSDQQLDQFFSVARAVSFFARAIDKYQHSDKESDKCIQTALRGLTEYVTSHHPCHHDANCQLSQGSSDPQPSGSNSSNDWTYGEAQLFARALEACGKNFGAIKKDFLPWKPVKSIIEYYYQGKEERPNSSHNNLGEPSTSMFSVVEPKKEIIDCDDLGDPHTSPDSGSSEEPDNSIPEERIDEPSDDKQDDETGSHPNLPTTSLTAEVLEVKPMKAKPVTATLDLETSPLGSLQFYFHGKLVLKLNAQQHANTNGHQCQWVPSVDTPKVPFPCSAGQCKKKYVELYEDSRTVDGADLQEVEGGFSKKAKLKNDFAIVWPEESFGAPVHSPQSPALSEWRHRGRATPYDGTLSPFSGVCSDPSDSDISPLQSPTDKCSTKKILGPSELCSVKDDLCIKPEPVSPCSHPTHNIEGRKIPFYSSCHKQDPQKFNTLEEVKRPFSQVNCTDVSQQIECSVPISYPSMLKTVLEATDKTGYKHLLSSSYKLSNVKQEVPLDLSIKSSQPSVNKLHAVETNCQTDGKLNSCSHKQKQLSSSWPDETLSHPPGLNTRRSSKQIKRSSFSSPDVRPSVSSFKSLTSSPVGGISKQHSADNKWTSINAKPLTLPPVISSATNTLHPGSATWTSFSANPDILSPTHTWEKSDATVASLTSYQEQHSGLTYLSMPIYNMAYPYPANCVENVNGVSASTEVNYFSQKNSPIKLKNMEDVTSSPSSKEKEIPAWVTNRPTDLTSYIPLYPHIYSYYPYAYNLSQVSWNGAVTPTEASDAPLDLSSPPSDNIPCSPKQKETIVGSSRAVQVPKDETDKDELSKLSNKSDDFSPSVSLGF
